# How does data structure impact cell-cell similarity? Evaluating the influence of structural properties on proximity metric performance in single cell RNA-seq data

**DOI:** 10.1101/2022.04.21.489121

**Authors:** Ebony Rose Watson, Ariane Mora, Atefeh Taherian Fard, Jessica Cara Mar

## Abstract

Accurately identifying cell populations is paramount to the quality of downstream analyses and overall interpretations of single-cell RNA-seq (scRNA-seq) datasets but remains a challenge. The quality of single-cell clustering depends on the proximity metric used to generate cell-to-cell distances. Accordingly, proximity metrics have been benchmarked for scRNA-seq clustering, typically with results averaged across datasets to identify a highest performing metric. However, the ‘best-performing’ metric varies between studies, with the performance differing significantly between datasets. This suggests that the unique structural properties of a scRNA-seq dataset, specific to the biological system under study, has a substantial impact on proximity metric performance. Previous benchmarking studies have omitted to factor the structural properties into their evaluations. To address this gap, we developed a framework for the in-depth evaluation of the performance of 17 proximity metrics with respect to core structural properties of scRNA-seq data, including sparsity, dimensionality, cell population distribution and rarity. We find that clustering performance can be improved substantially by the selection of an appropriate proximity metric and neighbourhood size for the structural properties of a dataset, in addition to performing suitable pre-processing and dimensionality reduction. Furthermore, popular metrics such as Euclidean and Manhattan distance performed poorly in comparison to several lessor applied metrics, suggesting the default metric for many scRNA-seq methods should be re-evaluated. Our findings highlight the critical nature of tailoring scRNA-seq analyses pipelines to the system under study and provide practical guidance for researchers looking to optimise cell similarity search for the structural properties of their own data.

## Introduction

Single-cell RNA-sequencing (scRNA-seq) enables investigation into the properties and heterogeneity of the individual cells within complex samples. The transcriptional profiles defining the current state of individual cells can be studied at high-resolution to identify signature genes, and patterns of expression which denote specific cellular processes [1], states [2], and types [3]. Accordingly, unsupervised clustering algorithms have become a popular approach in scRNA-seq for the identification of cell-types in an unbiased manner [4–6]. These algorithms partition cells into distinct clusters on the basis of cell-cell distances using a proximity metric, such as Euclidean distance.

Obtaining accuracy in clustering is hampered by structural properties of the scRNA-seq data. For example, there is an increased rate of dropouts in scRNA-seq compared to their bulk level counterparts, resulting in extremely sparse datasets and high levels of noise [7]. The capacity to measure thousands of features per cell, at scales of thousands to millions of cells, has led to increasingly high-dimensional data spaces. These spaces come with unique properties and limitations, often referred to as ‘the curse of dimensionality’ where the feature space is vastly larger than the sample space [8]. Furthermore, common clustering algorithms for scRNA-seq are based on the assumption that a dataset is composed of discrete and mutually exclusive groups of cells [4]. While these discretely-structured datasets do exist, such as tissue atlases which represent biological data of terminally differentiated cell types [9–11], datasets of continuous structure are also common. Continuously-structured datasets are composed of contiguous groupings of cells which experience multifaceted gradients of gene expression, encompassing dynamic processes such as embryonic development [12,13] and cell differentiation [14,15].

Knowledge of the biological system, and thus discrete or continuous structure of a dataset, is important as it has been shown to significantly influence the performance of certain scRNA-seq methods [4,16]. Heiser & Lau [17] identified that a dataset’s structural distribution is the primary determinant of dimensionality reduction performance, finding that preservation of structure is worse in discretely structured datasets than in continuous ones. The assumption of discrete, discernible cell-types in scRNA-seq clustering also poses challenges for identification of rare-cell populations, where rare cells may only differ from more abundant, stable cell populations by a small number of expressed genes [18–20]. Despite their low abundance, rare cell populations are of critical importance as they often represent highly specialised cell states or sub-types, and therefore provide valuable insights into processes such as differentiation, migration, metabolism and cancer [21–24]. It is also thought that all disease originates at the level of a single cell and therefore capturing rare cells accurately is critical for scRNA-seq analysis in clinical applications [25].

Intense efforts have been made to produce new clustering algorithms and benchmark the performance of existing methods to overcome the aforementioned limitations [4,26,27]. Some comparative studies specifically address data properties such as sparsity [28], dimensionality [29], rare-cell populations [30–34] and data structure [4,28,33]. Previous studies evaluating proximity metrics produce varied recommendations and lacked key design considerations. For example, Skinnider *et al*. [35] recommends proportionality-based metrics, whilst Kim *et al*. [36] recommends correlation-based metrics, specifically Pearson. However, Sanchez-Taltavull *et al*. [37] recommend Bayesian correlation over Pearson. Several other studies have proposed novel and scRNA-seq specific proximity metrics after observing variable performance of traditional proximity metrics [38–40]. Despite the inconclusive findings of previous works with respect to specific proximity metrics, they are largely in agreement that proximity metric performance is highly dataset-dependent [4,36,38,41,42]. This conclusion remains unworkable however, as the specific structural properties of the scRNA-seq datasets included in these evaluations are rarely addressed in detail or evaluated in a systematic manner.

Consequently, our study aims to address the important question of how the properties of scRNA-seq datasets influence the performance of proximity metrics in scRNA-seq cell clustering. To the best of our knowledge, such an investigation has yet to be performed and may be a reason why previous attempts that have been more limited, have been unable to yield actionable conclusions. Our study evaluates the clustering performance of 17 proximity metrics in the presence of a Continuous and Discrete data structure, under varying levels of cell-rarity, sparsity, and dimensionality that reflect the variability of real scRNA-seq data. Our findings demonstrate that taking the structural properties of the individual dataset into consideration when planning and executing an analysis pipeline leads to substantial improvements in performance of proximity metrics in scRNA-seq clustering. We believe similar performance gains may be possible in other analyses steps that use proximity metrics, such as dimensionality reduction and trajectory inference. Consequently, we provide readers with practical guidelines for selecting a preferred proximity metric and neighbourhood size with respect to the structural properties of their own datasets. We also provide our evaluation framework as a python package, *scProximitE*, to allow users to evaluate the performance of proximity metrics for their own datasets and structural properties of interest.

## Methods

### scRNA-seq Data Collection

The Discrete dataset was sourced from the CellSIUS benchmarking dataset [43] from Wegmann *et al*. [30] which profiled eight human cell lines. We used two subsets to evaluate how cell-population proportions influenced proximity metric performance (Figure 1). The first subset, Discrete Abundant, contains predominantly abundant cell populations, with seven cell-lines at proportions of low (5.4%) to high (32%) abundance, and one moderately rare population (2%). In contrast, the second subset, Discrete Rare, comprises of 6 rare cell-populations (0.08%-3.14%), and two highly abundant cell-populations at 40.15% and 50.21%.

**Figure 1:**
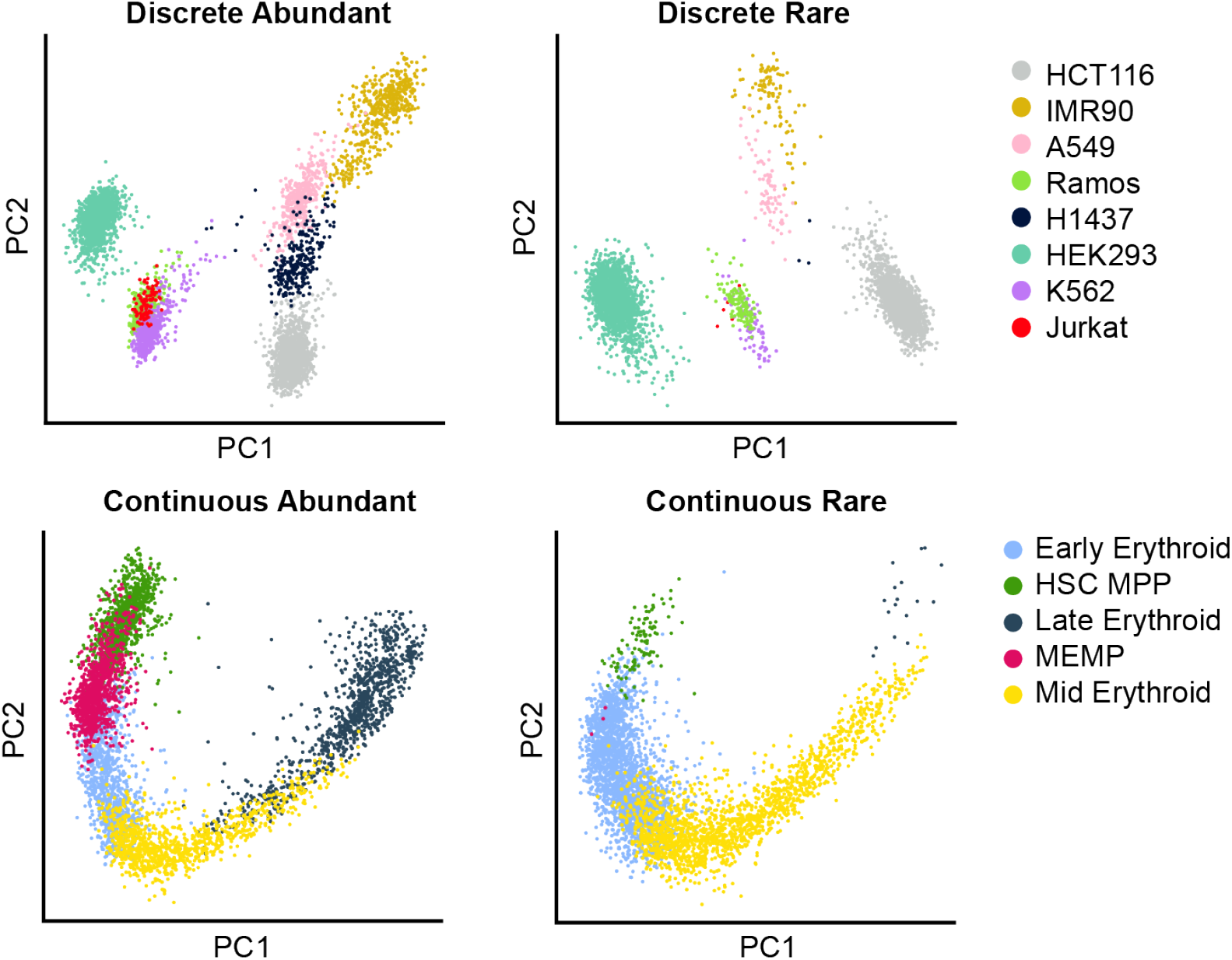
Principle Component Analysis (PCA) of the Discrete (top) and Continuous (bottom) structured scRNA-seq datasets, from the CellSIUS and Fetal Liver Haematopoiesis datasets respectively (see Methods). Datasets were subsampled to produce the Abundant datasets (left), representing data with only minor cell type imbalance, and Rare datasets (right), representing data with cell type proportion imbalances due to the presence of multiple rare cell populations.

The Continuous datasets consist of five erythrocyte differentiation cell-types from the Fetal Liver Haematopoiesis dataset (FLH) [44] produced in Popescu *et al*. [45] (Figure 1). As with the Discrete data, an Abundant and Rare subset were produced from the original FLH dataset. All five cell populations were present at high proportions (20%) in the Continuous Abundant dataset, whilst the Continuous Rare subset consisted of three rare cell-types present at proportions between 0.075% and 2.5%, and two highly abundant populations at 42% and 55%.

Both datasets were produced using 10x Genomics Chromium Single Cell (3’ library kit V2) and the provided cell-type annotations were used as the ground truth when evaluating clustering performance. For CellSIUS, single-cell sequencing was performed on batches of 2-3 cell-lines at a time, whilst bulk-sequencing was also performed for each cell line individually [30]. Cell-type annotations were generated by correlating single-cell profiles to the bulk profiles. For the FLH dataset, cell annotation was performed manually and validated through imaging mass cytometry, flow cytometry and cellular morphology [45]. The cell number and proportion for individual cell-types within each dataset are provided in Supplementary Table 1.

### scRNA-seq Data Simulations

Simulated datasets are used to evaluate how structural properties influence proximity metric performance, including, sparsity and cell-population imbalance. Simulated datasets follow a topology of four differentiation trajectories of equal length (3 branches), which diverge from a single origin state (1 branch). Each branch length is 50 pseudo-time units and represents a cell population in a continuous differentiation process. The PROSST package (v1.2.0) [46] was used to simulate 10 cells at each pseudo-time unit (6500 cells total) and 5000 genes from a negative binomial distribution. For each gene *g,* the variance parameters were sampled from *α_g_* ~ e^x^, x ∈ N (log_e_ (0.2), 1.5) and β*g* ~ e^x^, x ∈ N (log_e_ (1), 1.5), respectively. This simulated dataset in its original form was used to represent the Continuous Abundant Simulated dataset, whilst a subset containing only the origin state and the endmost population from each differentiation path was used to represent the Discrete Abundant Simulated dataset (Supplementary Figure 1, 2).

To further explore the influence of imbalanced cell-type proportions on metric performance, two structural subclasses, Moderately-Rare and Ultra-Rare, were created using Continuous Abundant Simulated and Discrete Abundant Simulated datasets. For the Moderately-Rare dataset, multiple cell-types are present at proportions *p* where 1% < *p* <5% whilst the Ultra-Rare datasets contain multiple cell-types where *p* <1%. The cell number and proportion for individual cell-types within each dataset are provided in Supplementary Table 2. The final structural property of interest in the study is dataset sparsity. Starting at 46-50% sparsity, two additional levels of moderate (68-71%) and high (89-90%) sparsity were produced for each of the six datasets by adding zeros using a Gaussian distribution (Supplementary Table 3).

### scRNA-seq Data Quality Control and Normalisation

The raw count matrices were filtered to remove i) cells with non-zero gene expression for <200 genes, ii) cells with >10% of their total counts arising from mitochondrial genes and iii) genes expressed in <10% of cells. The resulting cell and gene numbers for each dataset post-processing are in Supplementary Table 3 for simulated data, and Supplementary Table 4 for the CellSIUS and FLH datasets. Gene expression measurements for each cell were normalised by total expression and multiplied by a scale factor of 10,000, then log-transformed using natural logarithm, adding a pseudo count of one. All data processing steps, including filtering, normalisation, and identification of highly variable genes, were performed using the Scanpy package (v1.8.2) [47].

### Proximity Metrics

A total of 17 proximity metrics with a diverse range of properties were selected for evaluation. All metrics were computed in the form of dissimilarities, details regarding implementation are provided in Supplementary Table 5. For metrics implemented in R (v4.1.1) [48], Anndata [49] objects were converted to Seurat [50] objects using the sceasy package (v0.06) [51]. The formula for each proximity metric is available via the documentation of the relevant package. True distance metrics are dissimilarities which satisfy the four key properties of symmetry, reflexivity, non-negativity, and the triangle inequality. In this study, we included Euclidean, Manhattan, Canberra, Chebyshev, and Hamming distances. As the remaining 12 proximity measures do not strictly satisfy all four properties of a distance metric, we refer to all as “proximity metrics” herein for simplicity.

The proximity metrics, Hamming, Yule, Kulsinski and Jaccards Index, are computed on binary vectors. To generate the binarised count matrices for input, genes with ≥1 expression count were converted to 1, and genes with zero expression remained 0. All other metrics used normalised expression data as input.

Several of the evaluated dissimilarities are derived from correlations: Pearson, Spearman, Kendall, and Weighted-Rank correlations. Further selections included Bray-Curtis, a measure of compositional dissimilarity between two different samples, and Cosine, which measures the cosine of the angle between two vectors in the multi-dimensional space. In addition to commonly applied metrics, several recently proposed scRNA-seq metrics were also included. Given the sparse nature of scRNA-seq data, we evaluated the Zero-Inflated Kendall correlation (ZI-Kendall), an adaptation of Kendall’s tau for zero-inflated continuous data. Additionally, we evaluated Optimal Transport (OT) distance with entropic regularization, given its positive results for scRNA-seq clustering [52].As scRNA-seq data is relative rather than absolute, a proportionality-based metric called Phi was included, which Skinnider *et al.* [35] found to perform well in scRNA-seq clustering.

### Evaluation of Proximity Metric Performance

A key aspect of our study was to investigate how the presence and proportion of rare-cell types influence proximity metric performance, as such, we selected the Pair Sets Index (PSI) for evaluation of clustering performance [53], using the genieclust implementation (v1.0.0) [54]. PSI is a cluster validation metric based on pair-set matching and adjusted for chance, with a range of 0-1, where 0 indicates random partitioning whilst 1.0 represents perfect labelling with respect to ground truth annotations. Alternative evaluation measures were tested, including the Adjusted Rand Index (ARI) [55] and Adjusted Mutual Information (AMI) [56] (Figure 3), for which the Scikit-learn (v1.0.1) [57] implementation was used.

### Performance Evaluation Framework

The single-cell RNA-seq datasets representing the four structural classes of interest (Discrete Abundant, Discrete Rare, Continuous Abundant, Continuous Rare) (Figure 1) were first pre-processed and then used as input to calculate the distance matrix for each of the 17 proximity metrics (Figure 2). For each distance matrix, *k*-nearest-neighbour (KNN) graphs were computed using Scikit-learn library (v1.0.1) [57] in Python (3.8.11) [58], with the metric parameter set to ‘precomputed’. The KNN graphs are connectivity matrices, where each cell is connected to its *k* closest cells, as determined by the input distance matrix. To account for varying degrees of local structure, KNN graphs were constructed for each proximity metric at four different neighbourhood sizes: 3, 10, 30, 50. The resulting connectivity matrices are provided as input to the Scanpy implementation of the Leiden algorithm [47,59].

**Figure 2:**
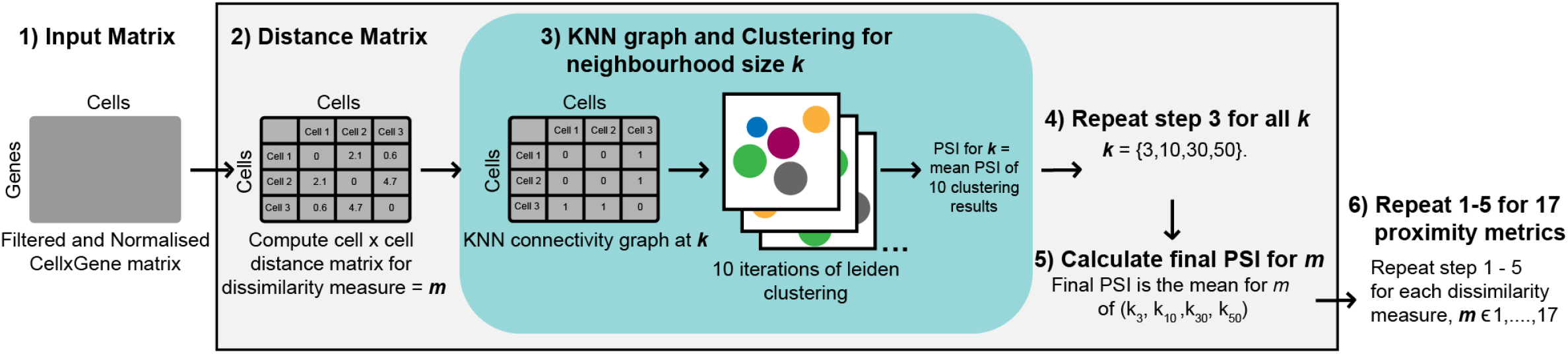
Evaluation framework for the assessment of the clustering performance of different proximity metrics. **1)** Processed scRNA-seq cell x gene matrices are the input to the framework. **2)** A distance matrix is calculated on the cell x gene matrix using a proximity metric. **3)** For each distance matrix, a KNN connectivity graphs is generated, on which Leiden clustering is performed 10 times using randomly generated initialisation values. The PSI is calculated for each clustering output, and the mean of the 10 results is taken as the PSI of the proximity metric at the *k* value. **4)** Step 3 is repeated for the proximity metric at four different neighbourhood size values (k=3, 10, 30, 50). **5)** The mean PSI from all neighbourhood sizes is calculated as the overall PSI value for the proximity metric. **6)** This pipeline is completed for each of the 17 proximity metrics included in this study.

The Leiden algorithm identifies clusters as groups of cells that are more densely connected to one-another than to the cells outside of the group based on the KNN graph [59]. Leiden is an unsupervised method with a resolution parameter that can be tuned to influence the number of communities detected. To accomplish accurate benchmarking, the resolution parameter was adjusted automatically until the number of clusters known to be present in the ground-truth cell-type annotations were returned, or until 1000 iterations had been attempted. To account for initialisation bias, 10 random seed values were generated, and the Leiden clustering was repeated with each seed for each connectivity graph.

The performance of the individual clustering outputs for each connectivity graph was compared to ground-truth cell annotations and quantified using PSI. The mean PSI across the clustering outputs was used to evaluate the neighbourhood size, *k*. Lastly, a mean PSI value was computed across the four neighbourhood sizes to summarize a proximity metric’s performance on a dataset.

## Results

We aimed to evaluate the performance of 17 proximity metrics in the presence of four types of scRNA-seq data structures: Discrete Abundant, Discrete Rare, Continuous Abundant, and Continuous Rare. A representative dataset was constructed for the Discrete structure from the CellSIUS benchmarking dataset [30,43] and included cells from eight distinct human cell lines (Figure 1). The Continuous structure category was represented by a subset of five erythrocyte differentiation cell types from the Fetal Liver Haematopoiesis dataset of the Developmental Human Cell Atlas [44,45].

Within the Continuous and Discrete datasets, a subclass was defined to reflect the balance of cell type proportions (Figure 1). An Abundant data structure is one where the majority of cell populations are present at a relatively high level, specifically, the majority of cell populations are represented at a proportion of ≥ 5% of the total cell number. In contrast, a Rare data structure is one where a majority of cell populations are represented at proportions of <5% (Supplementary Table 1).

We developed an evaluation framework to assess how the selected metrics performed with respect to properties relevant to scRNA-seq data (Figure 2). Specifically, the data properties were 1) data structure (Continuous or Discrete), 2) sparsity, and 3) cell rarity, 4) dimensionality, and 5) neighbourhood density. Comparisons to the ground-truth cell annotations were assessed using the Pair Set Index (PSI) for each metric.

Clustering results were obtained for ≥ 9 of the 10 repeats for all proximity metrics from the CellSIUS and FLH datasets, apart from Kulsinski at neighbourhood size 50 in the Discrete Abundant Dataset, which obtained 7. For the simulation datasets, ≥ 7 of the 10 repeats were returned, with two exceptions. For Cosine in the high sparsity Discrete Rare dataset at *k* = 100 only 4 results were obtained, and for Canberra in the high sparsity Continuous Rare dataset at *k* = 3, where only 1 result was returned.

We selected PSI to evaluate the clustering performance as incorrect clustering of rare and abundant cell-populations effects the final score equally [53]. Several popular evaluation metrics were considered for inclusion, including the Adjusted Rand Index (ARI) and Adjusted Mutual Information (AMI), however, we found the clustering score was dominated by the performance on Abundant populations, with little influence from Rare populations. For example, ARI and AMI scored a clustering output as near perfect on the Discrete Rare dataset (0.97, 0.91 respectively) despite 6 of the 8 cell-types being incorrectly clustered (Figure 3). Almost equivalent scores (ARI=0.98, AMI=0.96) were achieved by a clustering output where 6 of the 8 cell-types were accurately identified, showing the inability of these metrics to effectively distinguish clustering quality on datasets with substantial cluster-size imbalances. In comparison, PSI scored the second clustering result substantially higher (0.85) than the first (0.31). PSI has also been shown to be less sensitive to other clustering parameters such as the number clusters and degree of cluster overlap [53].

**Figure 3:**
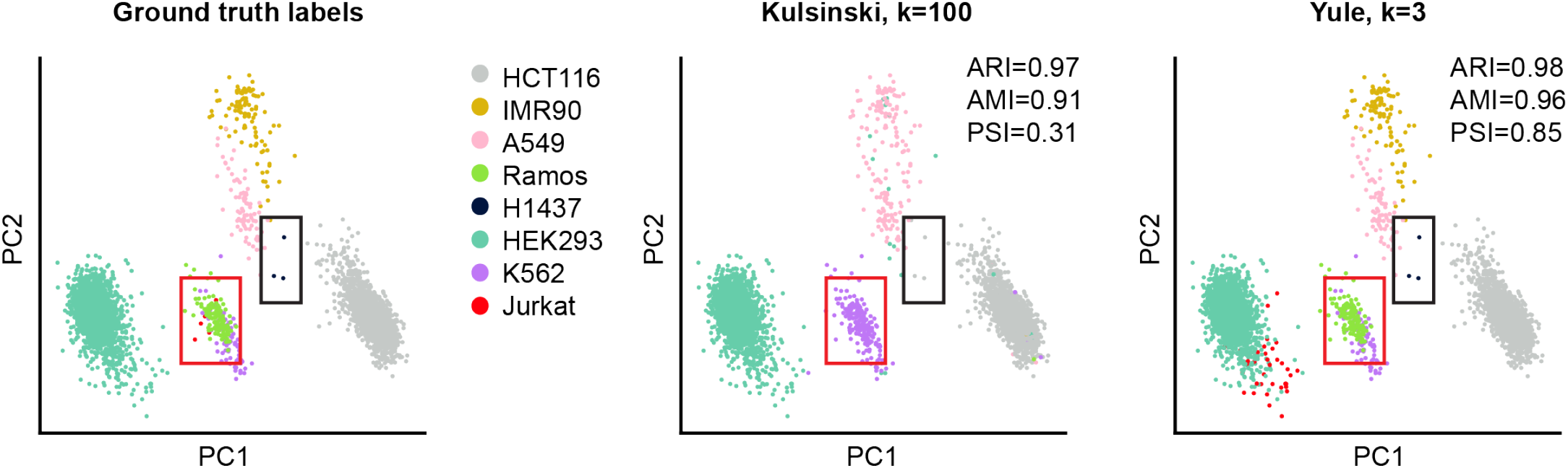
PCA in A) depicts the ground truth cell annotations for the Discrete Rare dataset derived from the CellSIUS benchmarking dataset. PCA in B) depicts the clustering results for the Discrete Rare data with the Kulsinski metric at a neighbourhood size of *k*=100. PCA in C) depicts clustering results for the Discrete Rare data with Yule at a neighbourhood size of *k*=3. Boxes are included to emphasise the location of the rare cell types H1437 (navy) and Jurkat (red) in each plot.

### Clustering performance of proximity metrics is dependent on the intrinsic structure of scRNA-seq datasets

We find that the capacity of proximity metrics to identify similarities between cells correctly in scRNA-seq data varies significantly depending on the intrinsic structure of scRNA-seq data (Figure 4). On average, proximity metrics achieved higher clustering performance for the Discrete data structures than the Continuous ones (on average by 0.4 PSI) (Figure 4). Within these structures, greater performance was observed for Abundant datasets than for Rare (by an average of 0.34 PSI) (Figure 4). We found that the magnitude of differences in clustering performance was larger between dataset structures than between metrics evaluated within the same structure. For example, the standard deviation across all metrics within the Discrete Abundant structure was only 0.097, whilst the standard deviation for the Euclidean distance metric across the four data structures was 0.27 (Figure 4).

**Figure 4:**
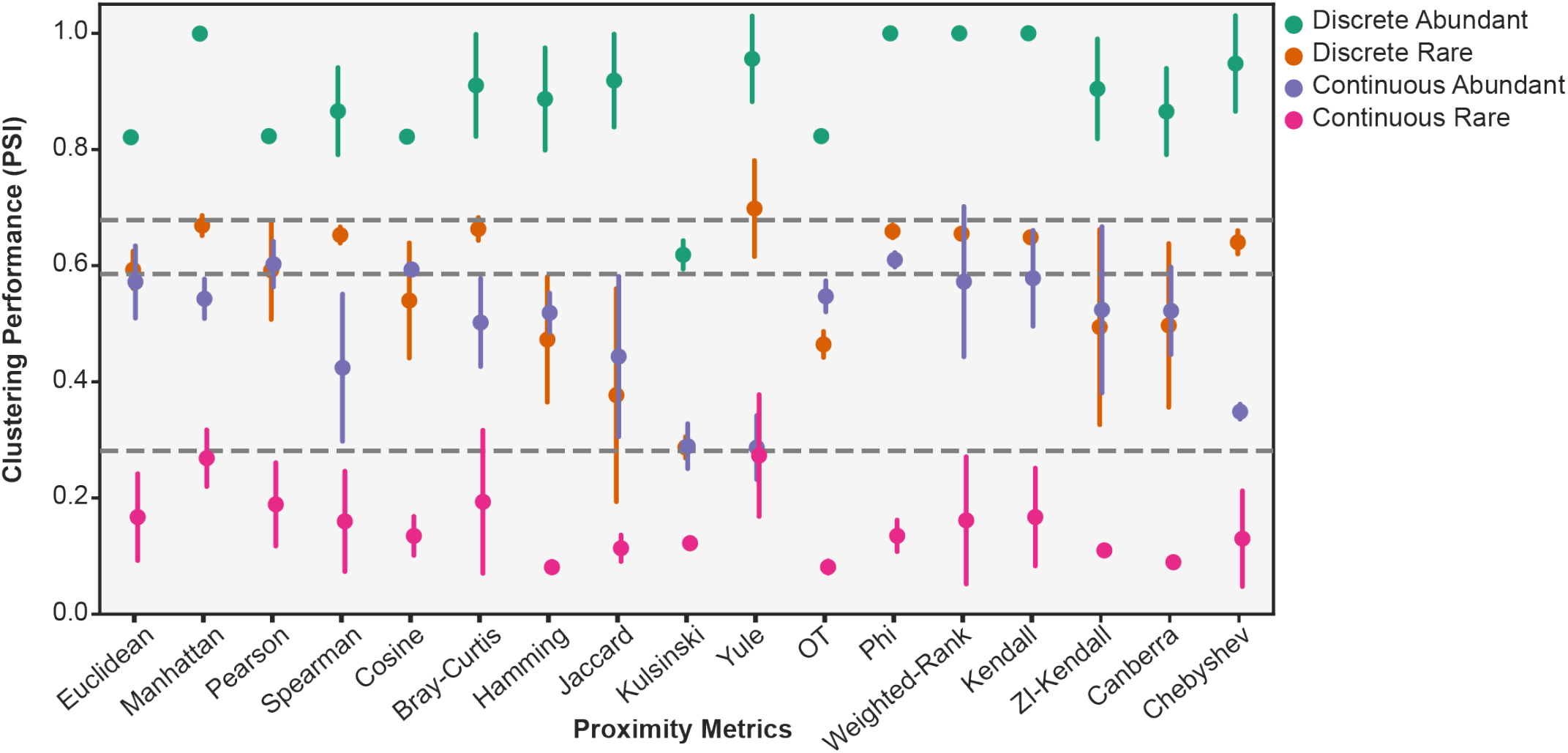
Clustering performance of proximity metrics for the scRNA-seq datasets representing the four classes of data structure: Discrete Abundant, Discrete Rare, Continuous Abundant, Continuous Rare. Points depict the mean Pair Sets Index (PSI) of clustering from neighbourhood sizes of *k* = (3,10,30,50), with error bars depicting the standard deviation. Horizontal lines depict (top to bottom) 75^th^, 50^th^ and 25^th^ percentiles. Error bars show one SD.

For the Discrete Abundant dataset, 16 out of 17 proximity metrics had good performance (PSI >= the 75^th^ percentile, Figure 4), whilst in the Discrete Rare dataset only a single proximity metric, Yule, performed at this level. Performance on the Continuous Abundant dataset was substantially lower, with only 3 metrics exceeding the 50^th^ percentile (Pearson Correlation, Cosine, Phi) (Figure 4). Continuous Rare datasets exhibited the poorest performance, with all metrics falling below the 25^th^ percentile. Similar trends are observed for simulated datasets (Supplementary Figure 3).

### Dimensionality Reduction Reliably Improves Clustering Performance of Proximity Metrics in Discretely structured datasets, but not Continuously structured

When working with high dimensional, noisy data like scRNA-seq data, it can be advantageous to perform dimensionality reduction, especially prior to clustering. However, the resulting changes to the data structure may cause some proximity metrics to work sub-optimally. To evaluate how dimensionality reduction affects performance of the 17 proximity metrics we reduced the dimensionality by selecting the 2000 most highly variable genes (HVG2000) and the 500 most highly variable genes (HVG500) and compared the performance to the full high-dimensional (HD) datasets. Metrics were considered invariant between any two levels of dimensionality if there was <0.05 change in the PSI score.

As expected, an improvement in performance between the HD dataset and at least one of the HVG datasets was observed for a range of proximity metrics in all structural classes (Supplementary Figure 4, 5). For the Discrete Abundant; Euclidean, Canberra, Hamming, Pearson, Spearman, Cosine and OT improved from <0.9 PSI to achieve near perfect clustering accuracy (>0.99 PSI) after the application of dimensionality reduction (Figure 5A, Supplementary Figure 5). A similar trend was observed for Euclidean, Canberra and Hamming in Discrete Rare, which ranked among the 5 highest performing metrics after dimensionality reduction to 500 HVG, despite their relatively poor performance in HD (0.47-0.59 PSI) (Supplementary Figure 5). This indicates dimensionality reduction is of particular benefit to the distance metrics commonly applied in scRNA-seq analysis. Conversely, despite experiencing the largest improvement in PSI (>0.28) due to dimensionality reduction for both Discrete structures, the binary metric Kulsinski remained the worst performing metric in these datasets (Figure 5A).

**Figure 5:**
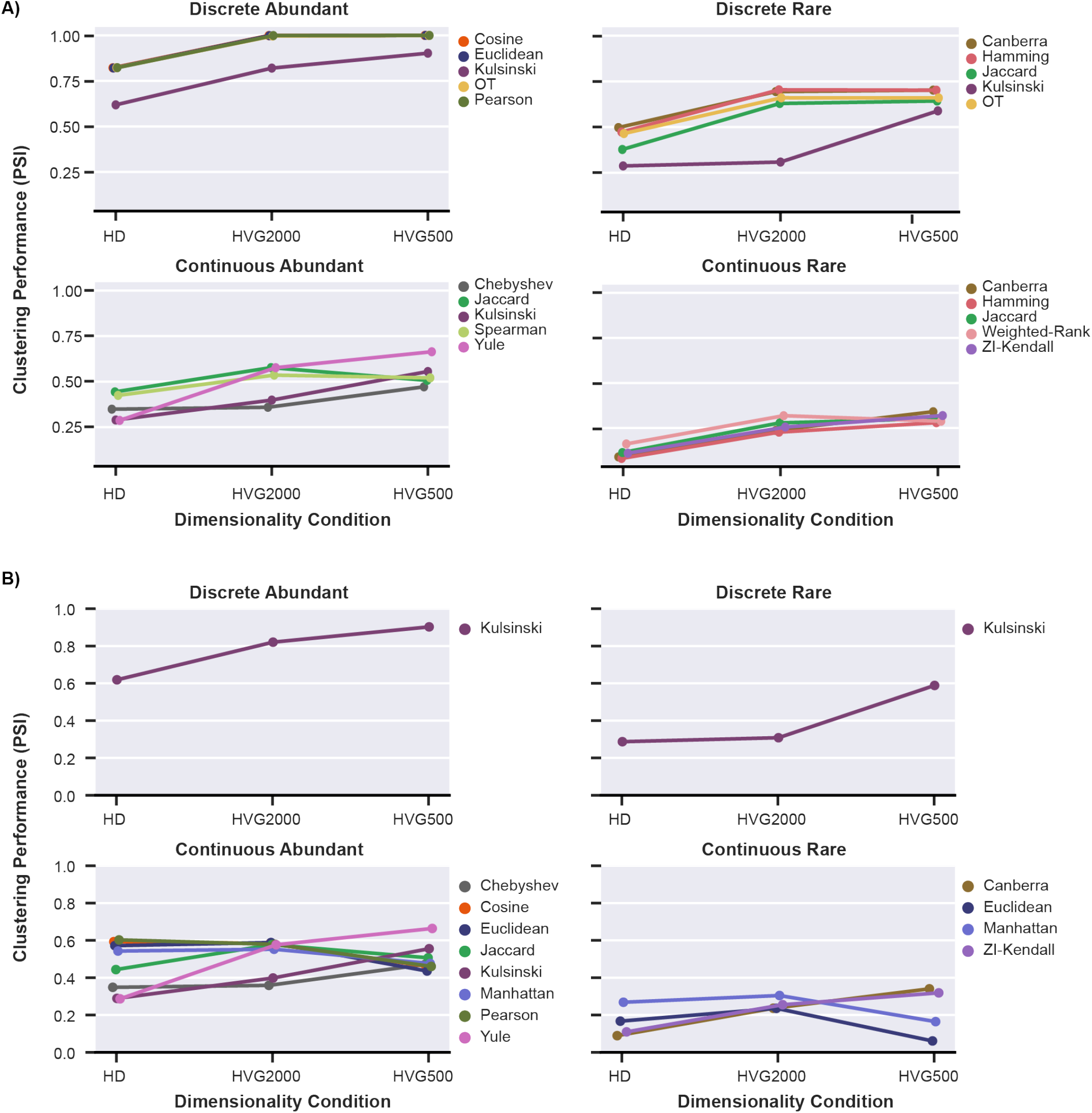
Clustering performance of **A)** the top five metrics for each structural condition after ranking for the greatest positive change between the high-dimensional (HD) dataset and either level of dimensionality reduction (HVG2000, HVG500), and **B)** metrics which showed >0.05 change PSI between the two levels of dimensionality reduction (HVG2000, HVG500), for each structural condition. Each point represents a PSI measurement of clustering performance from real scRNA-seq datasets that has been averaged across the neighbourhood sizes (*k* = 3,10, 30,50).

Despite substantial improvements in clustering performance due to dimensionality reduction, metrics in Discrete Rare data structures have lower PSI values (<0.71) than Discrete Abundant data structures. Similarly, when evaluating the Continuous data structures, the metrics with the largest improvement due to dimensionality reduction nevertheless had overall lower PSI values than the Discrete structure, PSI < 0.67 for Continuous Abundant, and <0.34 for Continuous Rare (Figure 5A). Accordingly, the trends of poorer clustering performance for scRNA-seq datasets with Continuous and/or Rare structure that are observed at HD largely remain after dimensionality reduction.

We use our findings to identify metrics with robust performance across HD, HVG2000 and HVG500. These metrics are characterized by a high level of performance as well as an invariant PSI across HD and HVG conditions and may be an attractive option when performing dimensionality reduction is not feasible. We defined a high-performance metric as one with a PSI at HD within 0.05 PSI of the maximum PSI achieved for either level of dimensionality reduction within the corresponding dataset. For the Discrete Abundant data structure, five metrics are identified as robust: Manhattan, Kendall, Phi and Weighted Rank, with a HD PSI >0.99, and Yule with a HD PSI >0.95. Consistent with this, Yule (0.7 PSI), Manhattan (0.67 PSI) and Phi (0.66 PSI) were also identified as robust metrics in the Discrete Rare dataset, along with Bray-Curtis (0.66 PSI) (Supplementary Figure 6). Of the few proximity metrics identified as invariant for the Continuous Abundant (3) and Continuous Rare (4) datasets (Supplementary Figure 6), none were classified as high performing, indicating that dimensionality reduction has a greater influence on datasets with continuous structure and therefore is likely to be a necessary step prior to clustering continuous datasets.

Given the limited number of metrics showing invariance to dimensionality on continuously structured data, we explored whether the extent of dimensionality reduction applied (HVG2000 vs HVG500) had an influence on proximity metric performance. Almost half of the proximity metrics (8/17) showed variable performance between the two HVG conditions in the Continuous Abundant data. Whilst Yule, Kulsinski and Chebyshev performed best at HVG500, the remaining five metrics, including Euclidean and Manhattan, showed poorer performance at HVG500 relative to HVG2000 (Figure 5B). For Continuous Rare, Canberra, Hamming and ZI-Kendall showed improvement from HVG2000 to HVG500, whilst as observed in the Continuous Abundant data, Euclidean and Manhattan showed the reverse (Figure 5B). In comparison, 16 of the 17 proximity metrics in the Discrete datasets exhibited robust clustering performance between 2000HVG and 500HVG, with the outlier being Kulsinski (Figure 5B). This suggests that in discretely structured data, equivalent information may be captured with 500 genes as with 2000 for most metrics, but also that further reduction beyond 2000HVG does not provide additional benefits such as greater reduction of noise. Conversely, for continuously structured data there may be a narrower parameter range at which the benefits of dimensionality reduction are balanced with the loss of relevant structural information.

### All Proximity Metrics are Sensitive to Increasing Rarity of Cell-Populations

The identification of rare-cell types in scRNA-seq remains challenging, as demonstrated by the consistently poor clustering performance achieved for our Rare datasets, regardless of structure or dimensionality. The rare subsets from CellSIUS and FLH datasets contained ‘rare’ cell types at proportions ranging between 0.08%-3.1% and 0.425%-2.5% respectively (Supplementary Table 1). To investigate if proximity metric performance is only impacted beyond a certain threshold of rarity, we used the PROSSTT package to simulate scRNA-seq datasets with Discrete and Continuous structure, for which we generated an Abundant (all populations >5%), Moderately-Rare (multiple populations at >1% to <5%), and Ultra-Rare dataset (multiple populations at <1%) (Supplementary Table 2). Datasets representing our moderate-sparsity condition (68-71% zero values) were used for the results included (Figure 6), however results for all sparsity levels are depicted in Supplementary Figure 7.

**Figure 6:**
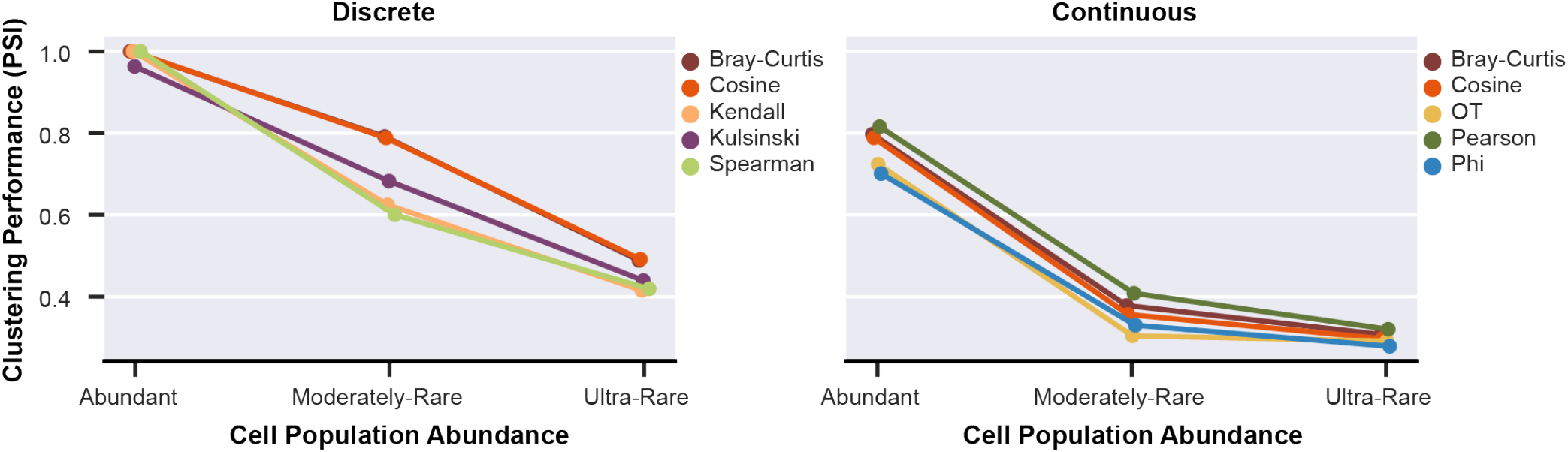
Clustering performance of the top five proximity metrics, as ranked by PSI on the Ultra-Rare simulated scRNA-seq dataset (moderate sparsity), for Discrete (left) and Continuous (right) structure. Points depict the mean Pair Sets Index (PSI) of clustering from neighbourhood sizes of *k* = (3,10,30,50), with error bars depicting standard deviation.

We observed a substantial reduction in clustering performance between the Abundant and Moderately-Rare datasets for Discrete (mean change in PSI = 0.29, SD=0.09) and Continuous (mean change in PSI = 0.24, SD=0.17) datasets, indicating that cells at proportions of ≥1% are sufficiently rare to challenge proximity metrics (Figure 6). Between the Discrete structure Moderately- and Ultra-Rare datasets, performance was further reduced by a mean of 0.23 (SD=0.07) across all metrics, with the maximum PSI achieved at 0.49. Notably, there was no significant difference in PSI from Moderately-Rare to Ultra-Rare datasets of Continuous structure (mean change in PSI = 0.04, SD = 0.03). This is unsurprising given that the metrics already displayed very poor performance for identifying Moderately-Rare cell types (≤0.41 PSI) (median PSI 0.28, SD = 0.08). Notably, whilst Bray-Curtis and Cosine were included in the top 5 performers for both Discrete and Continuous data structures based on PSI in Ultra-Rare datasets (Figure 6), no proximity metrics showed significantly greater robustness to the presence of increasingly rare cell populations.

Our findings suggest that there is not a threshold of ‘rarity’ (cell-population proportion) at which the performance of proximity metrics is suddenly impacted, but rather a continuing decline in performance for cell-populations of decreasing proportions relative to the total dataset. We show the proximity metrics’ capacity to capture structural information is particularly challenged in datasets comprised of cell-populations representing continuous processes and datasets containing rare cell-populations.

### Most metrics perform worse as sparsity increases, but under-utilised metrics show greater robustness

Sparsity is one of the greatest challenges when working with scRNA-seq data and hence it is important to evaluate performance against this structural property. Using our Abundant and Moderately-Rare (hereafter referred to as ‘Rare’) simulated scRNA-seq datasets, we adjusted sparsity to 3 different levels: low (the baseline sparsity of 46-50%), moderate (68-71%) and high (89-90%) by simulating additional dropout based on a Gaussian distribution (Supplementary Table 4). To interpret the results, we defined a metric as robust to sparsity if the change between PSI levels for different sparsity conditions was ≤0.05, sensitive if the change between PSI levels was ≥ the 75^th^ percentile for all metrics in that structural class, and moderately sensitive if between these thresholds (Supplementary Figure 8).

Similar to the influence of dimensionality reduction, proximity metrics are influenced by sparsity to a greater degree on continuously structured data than on discretely structured data. Encouragingly, a substantial number of proximity metrics demonstrated robust performance when sparsity was increased from low to moderate for the Discrete Abundant (11/17) and Rare (7/17) datasets (Figure 7). Conversely, no metrics were identified as robust for Continuous Abundant, and only two in Continuous Rare. Notably, Bray-Curtis and Pearson correlation were identified as robust metrics for the Discrete Abundant, Discrete Rare, and Continuous Rare structured datasets. Furthermore, Bray-Curtis, Cosine and Pearson correlation were consistently ranked among the top five metrics with the least sensitivity to sparsity for all structural conditions (Supplementary Figure 9). However, it should be noted, the maximum PSI for the Continuous Rare dataset with moderate sparsity was only 0.41, indicating that the clustering performance of even the best ranked proximity metrics was poor for this structure.

**Figure 7:**
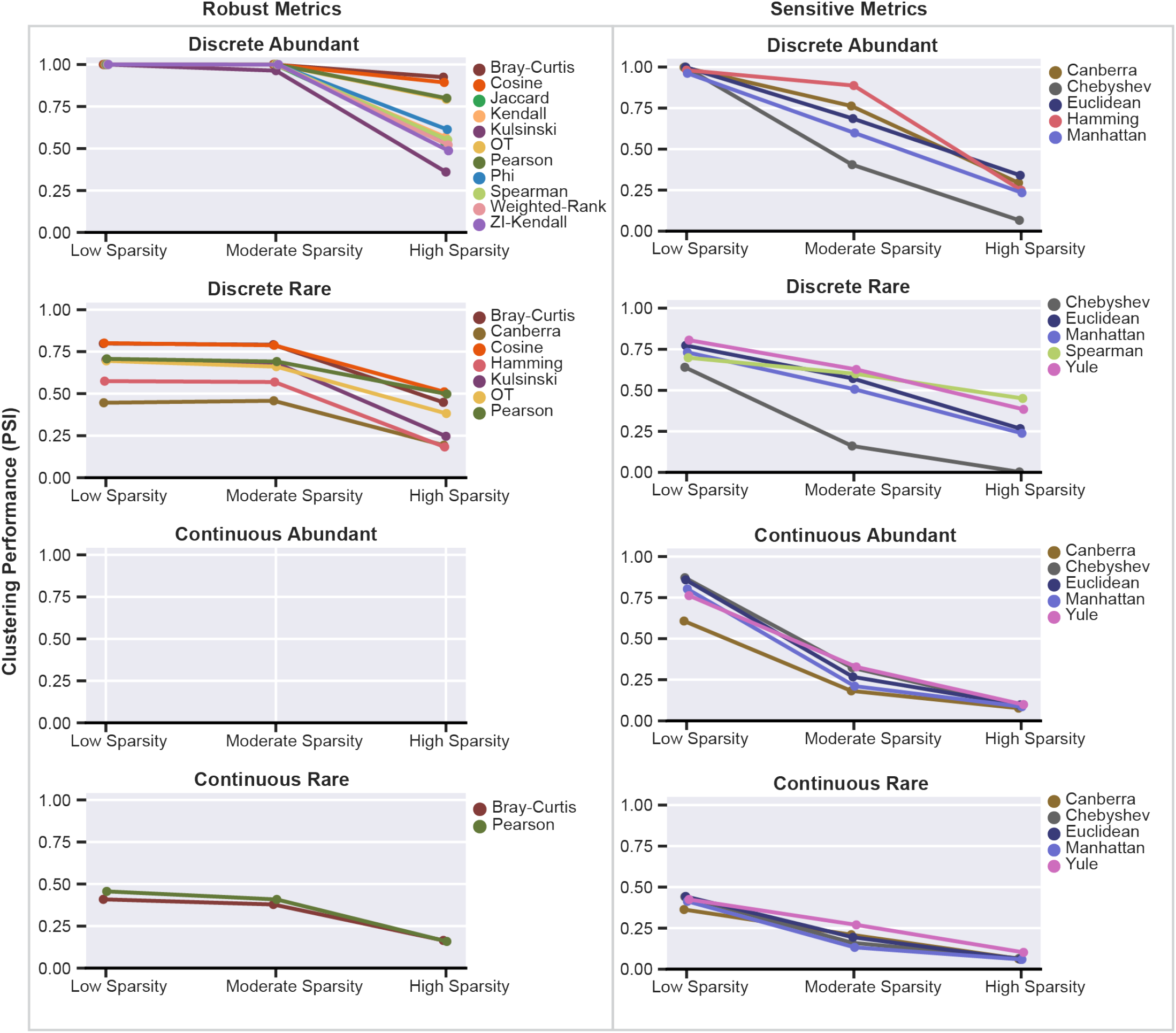
**Left** – Performance of proximity metrics identified as robust between low (50%) and moderate (70%) sparsity, given a threshold of ≤0.05 change in PSI. As no metrics met these criteria for the Continuous Abundant dataset, the panel is blank. **Right** – Performance of proximity metrics identified as sensitive between low and moderate sparsity, given a threshold of ≥75^th^ percentile change in PSI. Points depict mean PSI of clustering performance from simulated scRNA-seq datasets across neighbourhood sizes of *k* = (3,10, 30,50) and the four classes of structure, for each level of sparsity described.

Interestingly, performance of the true distance metrics were more sensitive to sparsity than the other proximity metrics, with Euclidean, Manhattan and Chebyshev being the only metrics that were sensitive to sparsity across all four conditions, whilst Canberra was sensitive in all data structure types except Discrete Rare (Figure 7). This result reflects that the true distance metrics share a fundamental property resulting in more sensitive performance in sparse, high-dimensional data. Yule, one of the proximity metrics based on Boolean vectorisation of gene expression data, was also identified as sensitive in three of the four structural conditions. Our results suggest that Bray-Curtis, Cosine and Pearson correlation may be the preferred metrics when analysing datasets with moderate levels of sparsity, as opposed to the more commonly applied Euclidean and Manhattan distance metrics.

Despite maintaining clustering performance at moderate sparsity, all ‘robust’ proximity metrics drop substantially in clustering performance when applied on the high sparsity data. Furthermore, at high sparsity the difference in performance between Abundant and Rare structures becomes equivalent in the Continuous dataset, with a maximum PSI of 0.21 (Supplementary Figure 10). This indicates that insufficient information is present in highly sparse scRNA-seq data to enable the discrimination of contiguous cell types, irrespective of cell-population abundance. The same trend is observed for the Discrete data, with the exception of Bray-Curtis, Cosine and Pearson correlation which provide good clustering performance for Abundant datasets (≥0.8 PSI). Consequently, reduction of dataset sparsity, whether via filtering, dimensionality reduction or imputation, is a key factor in optimising performance of proximity metrics on scRNA-seq data, with particular necessity for continuously structured data.

### Dataset Structure and Sparsity are key factors in clustering parameter optimisation

Clustering approaches based on KNN graphs, such as the Leiden algorithm, have grown popular in scRNA-seq analysis due to their speed and scalability. These methods are unsupervised, with a tuneable parameter, *k*, the number of nearest neighbours to identify for each cell. As the neighbourhood size, *k*, affects the number and size of clusters identified, selecting the appropriate *k* for a dataset is important. We investigated the impact of the neighbourhood size by varying *k* across five levels (3,10,30,50,100) and testing the performance of the metrics under each simulated data structure and sparsity condition. To identify metrics with the strongest performance across all neighbourhood sizes, we focused on those with a maximum PSI value across all neighbourhood sizes ≥75^th^ percentile (Figure 8).

**Figure 8:**
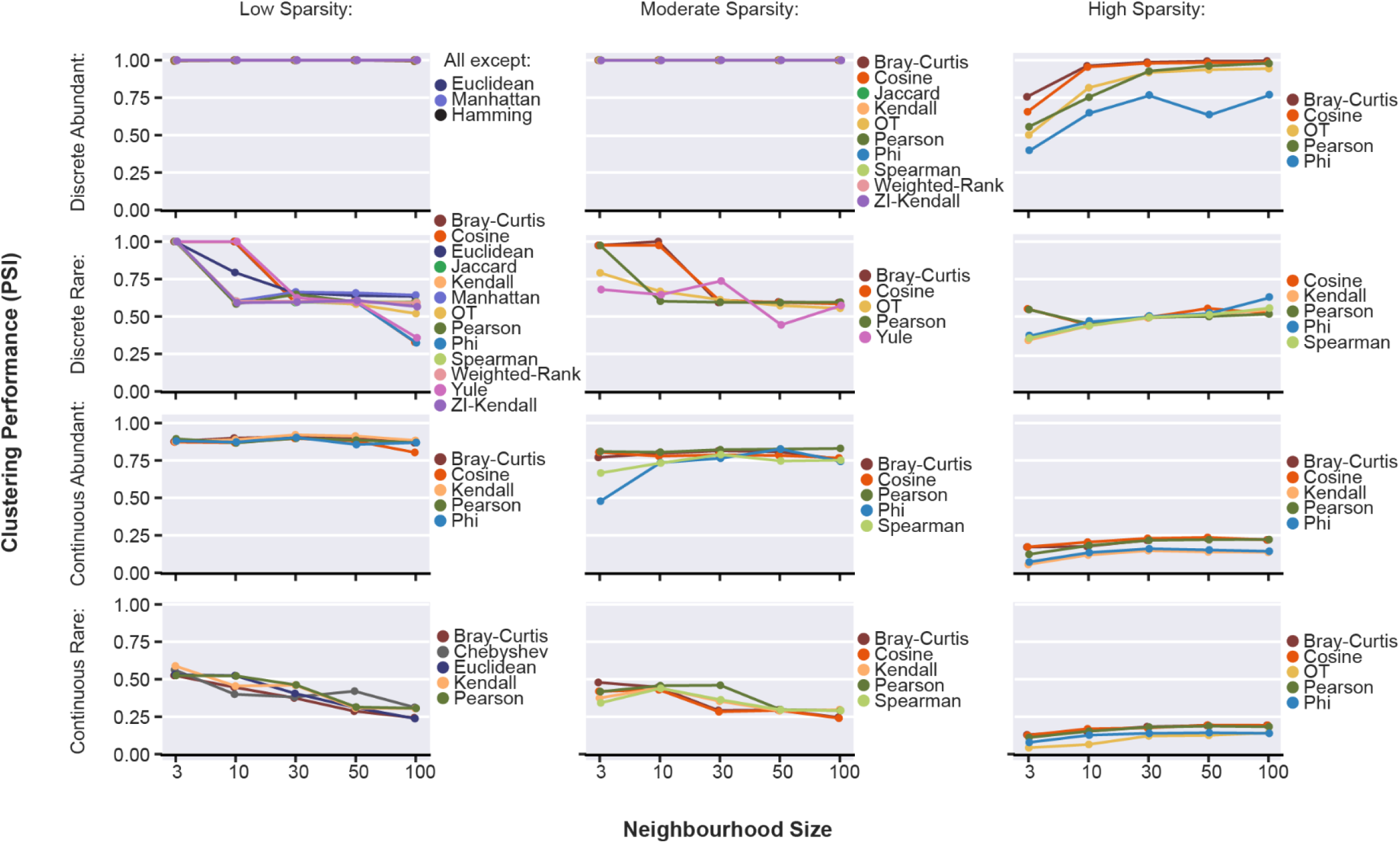
Clustering performance across the five neighbourhood size values for K-Nearest Neighbours, for low sparsity (Left), moderate sparsity (middle), and high sparsity (right) simulations. Proximity metrics are included if their maximum PSI achieved across all neighbourhood sizes is ≥75th percentile of the maximum performance in the relevant structural class.

At low sparsity, proximity metrics achieved greater performance in both Discrete and Continuous structure datasets containing Rare cell-populations at small neighbourhood sizes (3,10), whilst performance on Abundant datasets was invariant (Figure 8). In datasets of moderate sparsity, these trends are weaker as performance becomes more specific to the proximity metric. For example, Phi and Spearman correlation perform poorly for neighbourhood size of 3 in the Continuous Abundant data. However, these trends change noticeably at high sparsity, with proximity metrics showing increased performance at larger neighbourhood sizes (30,50,100) in the Discrete Abundant dataset. A similar but weaker trend is seen in the Discrete Rare dataset, except for Cosine and Correlation which continue to exhibit greatest clustering performance at a neighbourhood size of 3. In the Continuous datasets, performance is consistently very poor across metrics regardless of neighbourhood size (<0.25 PSI). The inconsistent relationship between neighbourhood size and clustering performance at high-sparsity further underlines the challenges associated with capturing structural information from highly sparse scRNA-seq datasets and supports our findings from our investigation of sparsity, reinforcing our recommendation to reduce dataset sparsity.

### Summary and Practical Recommendations

A summary of our findings are encapsulated in a flowchart to provide practical guidance on selection of appropriate proximity metrics based on the dataset (Figure 9). Recommendations have been derived from the set of metrics and neighbourhood sizes that were used during the clustering performance evaluation using the datasets in this study. Overall, the diverse nature of the proximity metrics evaluated in this study was exemplified in their differing responses to the structural properties investigated. For example, Cosine is the highest ranked metric for robustness to sparsity across all data-structures (Figure 10A) but responded inconsistently to dimensionality reduction (Figure 10B). In contrast, Manhattan distance performance was robust to changes in dimensionality (Figure 10B), but it is among the most sensitive metrics to even moderate sparsity (Figure 10A)

**Figure 9:**
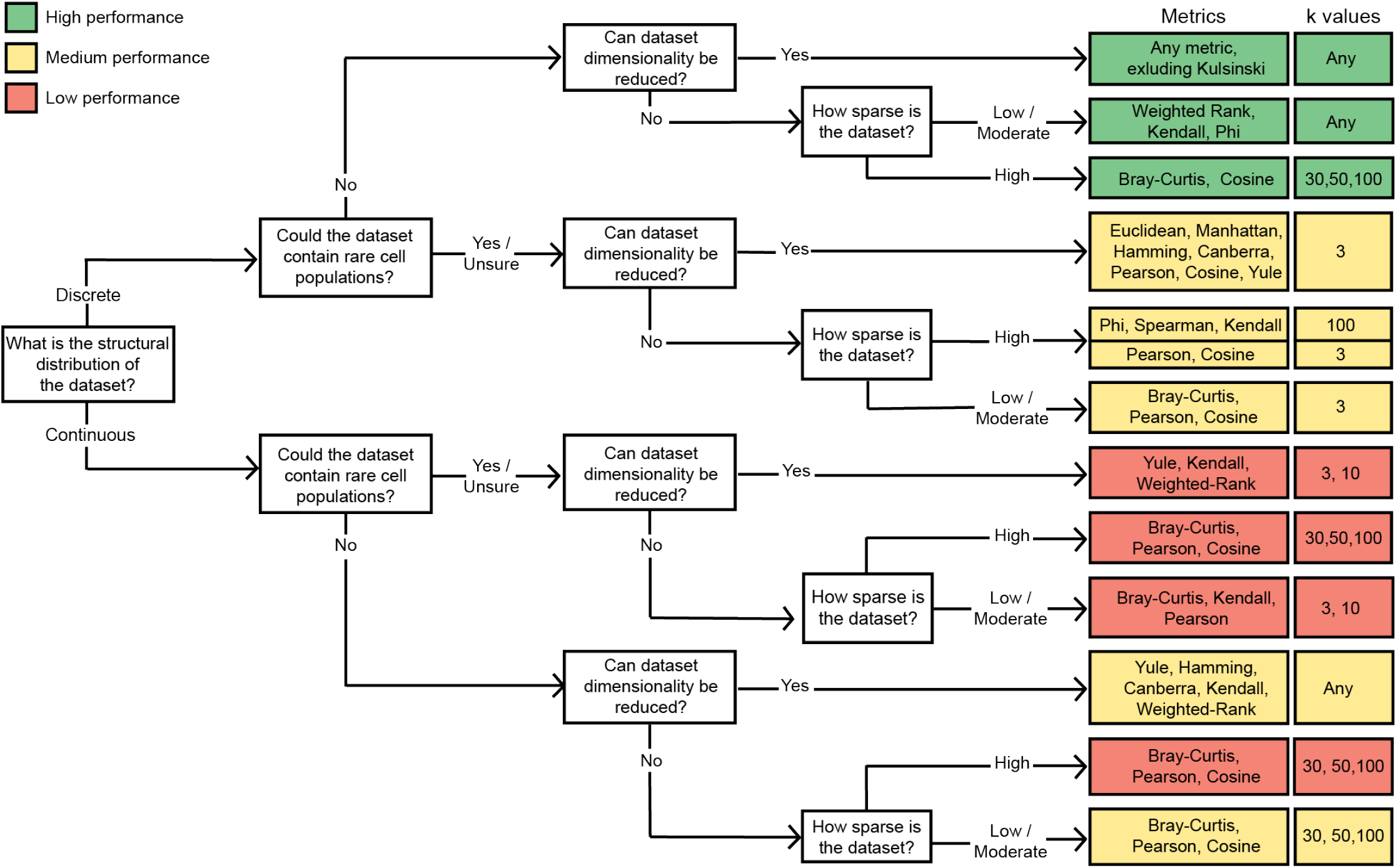
Flowchart for recommended metrics and neighbourhood sizes (*k*) given specific structural properties of a scRNA-seq dataset, based on results obtained from the evaluation framework on simulated and real scRNA-seq datasets. See Supplementary Table 6 for additional details.

**Figure 10:**
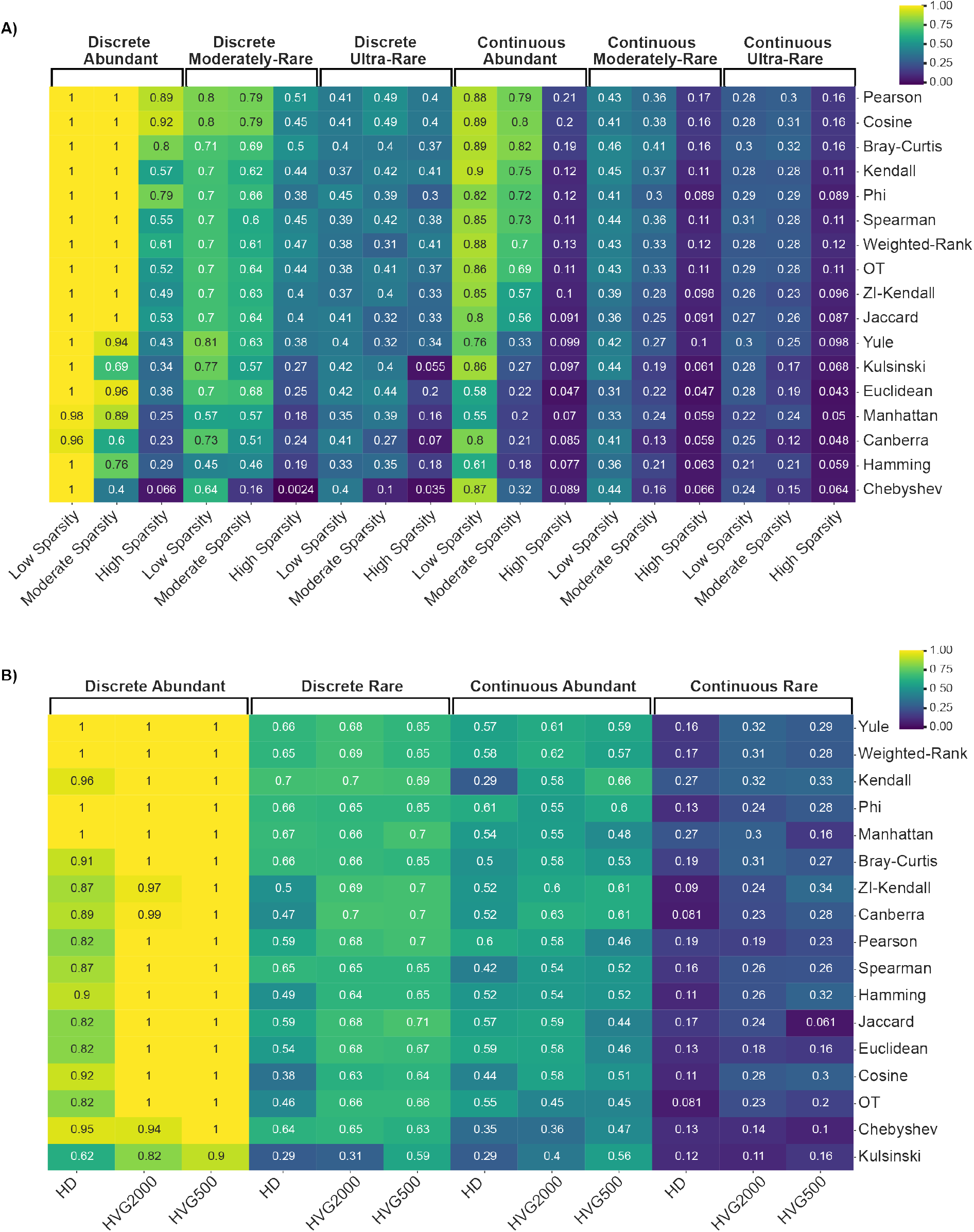
Proximity metric performance across real scRNA-seq datasets of varying structure and **A)** sparsity and **B)** dimensionality. Heatmap cells contain mean PSI obtained across neighbourhood sizes of 3,10,30,50, for each metric and dataset combination. Rows are ordered by mean PSI across datasets with strongest performance at the top.

Metrics which are recommended for ≥ 50% of the combinations of structural properties investigated include Pearson correlation (8/12), Cosine (8/12) and Bray-Curtis (7/12) (Figure 9), which showed the greatest robustness to dataset sparsity (Figure 10A). Kendall correlation (6/12), which was among the top five metrics for both dimensionality and sparsity (Figure 10), displayed a high degree of robustness relative to other metrics investigated. Conversely, despite its prominence in scRNA-seq analyses, Euclidean distance exhibited equivalent or lower performance than a range of less common metrics across the dataset structures, showing particular sensitivity to high-dimensionality (Figure 10B) and sparsity (Figure 10A). The adaptation of Kendall correlation for sparse data, ZI-Kendall, was also found to perform poorer than the original version under moderate and high sparsity conditions (Figure 10A).

In Discrete datasets composed of Abundant cell populations, application of dimensionality reduction ensured equally exceptional clustering performance (>99%) for all proximity metrics and neighbourhood sizes, excluding Kulsinski (Figure 10B). Similar performance could be achieved even in highly sparse (Figure 10A) or high-dimensional datasets (Figure 10B), given appropriate selection of proximity metric and neighbourhood size (Figure 9). For Discretely structured datasets containing multiple Rare cell-populations, Pearson correlation and Cosine were identified as top performing metrics at a neighbourhood size of 3 for all combinations of data properties. However, the greatest performance (0.71) could only be achieved in datasets which underwent dimensionality reduction or in which sparsity was low to moderate, highlighting the importance of the data processing step prior to clustering. The application of dimensionality reduction also expanded the pool of high performing metrics to include many of the distance metrics and the binary dissimilarity metric, Yule (Figure 10B).

For Continuous structures, data processing to reduce sparsity and dimensionality is necessary to optimise the performance of proximity metrics, regardless of cell population proportions. With application of dimensionality reduction, Yule, Kendall, and Weighted Rank were consistently among the top performing metrics for Abundant and Rare datasets, whereas the performance for a variety of other proximity metrics suffered upon greater dimensionality reduction (Figure 10B). In scenarios where datasets are unable to undergo dimensionality reduction, Pearson correlation, Bray-Curtis and Cosine were the highest performing metrics across all sparsity levels, although performance was significantly lower in the high sparsity datasets (Figure 10A). When figures were generated with PSI at 30 neighbours (the default value in Seurat), the top 5 ranked metrics remained the same for dimensionality, and top 4 metrics for sparsity, albeit re-ordered. This suggests our results may be relevant even without parameter tuning (Supplementary Figure 11).

## Discussion

Given the direct influence of cell clustering on downstream analysis in scRNA-seq data, evaluating the accuracy of clustering algorithms is an important area of research. Previous studies have recognised the effect of proximity metric choice when measuring cell-to-cell similarity on clustering performance [36,38]. However, variable performance is reported for proximity metrics between datasets, making the recommendation of a specific metric impossible [38]. In response, we developed a framework to evaluate 17 proximity metrics with respect to core structural properties of the scRNA-seq data, including sparsity, dimensionality, structure, and rarity. Our findings demonstrate that greater care should be taken to select and fine-tune methods to suit the structural properties of the individual biological system under study. Consequently, we have provided practical guidance for researchers to optimise their cell similarity search by investigating and acting on the structural properties of their own data.

Of the actions available, we identified reducing dataset sparsity as the most impactful factor for improving clustering performance (Figure 7). Sparsity reduction can be achieved via filtering and feature selection, but may require the application of scRNA-seq imputation methods [60,61]. Dimensionality reduction via selection of highly-variable genes also produced improvements in clustering performance for many proximity metrics (Figure 5). However, the variable results observed for continuously structured data indicate that the degree of dimensionality applied must be tuned appropriately to the dataset. How proximity metrics performance may be influenced by transformative dimensionality reduction approaches such as PCA [62], t-SNE [63,64] or UMAP [65,66] remains to be explored, but the influence of discrete and continuous data structure on these methods are reviewed in Heiser & Lau [17].

Selection of an appropriate neighbourhood size was essential for optimising performance of proximity metrics to accommodate cell-balance properties (Figure 8). Notably, the greatest performance for Rare datasets was obtained with neighbourhood sizes of 3 and 10, as opposed to the default values of 20 and 30 in Scanpy and Seurat, respectively. This finding illustrates the importance of tuning parameters for a given dataset based on knowledge of the underlying system, rather than relying on default settings [67]. The optimal parameter values for dimensionality reduction methods have been similarly shown to be a function of dataset-specific properties [17,68–70], and we expect that this extends to other aspects of scRNA-seq analysis.

We consistently identified cell-population structure to be one of the most influential properties, with substantially lower clustering performance for proximity metrics in datasets with continuously structured populations than discrete (Figure 4, Supplementary Figure 3). This has previously been identified as a shortcoming of clustering methods, and alternatives such as pseudo-time analysis [71] or soft clustering [72] have been proposed [4]. However, given that these recommended alternatives similarly rely on the calculation of cell-to-cell similarity, selection of an appropriate proximity metric is likely to be equally relevant. Additionally, performance was inferior in datasets with imbalanced cell-population proportions due to Rare cell populations, as compared to the Abundant datasets (Figure 4, Figure 6). Whilst we identified preferred dataset processing steps, proximity metrics and parameter values to improve performance on Rare datasets (Figure 9), we were unable to match the clustering performance of the Abundant datasets for either Discrete or Continuous structures.

It is worth highlighting that only by basing our evaluation framework around a performance score which is independent of cluster size, such as the PSI, could the true extent of this effect from rare cell-populations be revealed (Figure 3)[53]. It is likely that unsatisfactory clustering accuracy due to rare cell-populations is similarly present in other comparative evaluations, but largely masked by the use of evaluation scores such as ARI and AMI. For ARI and AMI cluster evaluations are size-dependent, thus the influence of misclassified rare cell-populations on the overall score is greatly diminished [53,73,74]. Given common approaches for data processing, normalisation, feature selection and clustering were used in the course of our study, these findings raise concerns regarding the current state of rare cell identification in scRNA-seq. A beneficial extension to our work would be to include specialised clustering methods developed for rare cell populations, such as GiniClust [31], scAIDE [32] or CellSIUS [30]. However, if researchers are unaware of the presence of rare cell types in their data, they will most likely fail to seek out such specialised methods. As such, there is a crucial need for greater integration of rare cell-type methods into popular scRNA-seq packages and standard analysis vignettes.

Euclidean distance is among the most commonly applied proximity metrics for cell-cell distance in scRNA-seq. Despite this, when evaluated for robustness to sparsity and high dimensionality in our datasets Euclidean, and the other true distance metrics evaluated, showed greater sensitivity relative to a range of lesser known proximity metrics (Figure 7, Supplementary Figure 6). These results were not entirely unexpected, as true distances metrics have been demonstrated to perform poorly as dimensionality and sparsity increase, leading to poorly defined nearest neighbours [75,76]. In line with this, we saw distance metrics perform considerably better with the appropriate level of dimensionality reduction, at times even achieving the maximum level of performance (Figure 5).

Our findings are supported by previous studies which have similarly identified Euclidean as a poorly performing proximity metric in scRNA-seq [35,36,38]. In Kim *et al.* [36] correlation-based metrics out-performed Euclidean distance for clustering, which was attributed to the sensitivity of the distance metrics to scaling and normalisation, whereas correlation-based metrics are invariant to these factors [36]. Interestingly, Pearson and Kendall correlations, along with another scale invariant metric, Cosine, were identified as the preferred metrics for the majority of structural conditions examined in our study. However, other scale-invariant metrics such as Spearman correlation did not show the same performance trends. Skinnider *et al.* [35] also found Euclidean performed poorly for a range of analysis tasks, including cell clustering, and suggested that as scRNA-seq only yields the relative abundance of gene expression within a cell rather than the absolute amount. Accordingly, they state that metrics of proportionality such as Phi and Rho are more suitable [35,77]. Whilst Phi performed moderately well in our evaluation, it was outperformed by Pearson, Kendall, and Cosine. However, another proportionality-based metric, Bray-Curtis, was identified as a preferred metric for more than half of the structural condition combinations evaluated.

Our results suggest that given the high dimensional and sparse nature of scRNA-seq data, the use of Euclidean distance as the default proximity metric should be re-evaluated. Several clustering methods that make use of proximity metrics aside from Euclidean have already been developed, have been shown to perform well for scRNA-seq data. For example, SC3 generates a consensus distance matrix derived from the Euclidean, Pearson and Spearman proximity metrics [78]. RaceID3 is a rare cell-type clustering method which allows the user to select from a range of distance and correlation-based metrics [79]. Other methods have instead developed entirely new metrics to measure cell-cell similarity, such as CIDER which recently proposed Inter-group Differential ExpRession (IDER) as a proximity metric for their new clustering pipeline [80].

While we aimed to design our study to be as comprehensive as possible, there are aspects of the framework which could be extended to evaluate additional factors, for example, expanding clustering methods to include approaches beyond graph-based clustering. However, similar results were obtained by Skinnider *et al.* [35] when they compared the clustering performance of proximity metrics with hierarchical and graph-based clustering approaches, suggesting that our results may hold for other methods. As with clustering, many scRNA-seq dimensionality reduction methods rely on the calculation of cell-cell similarity with a proximity metric. To minimise the influence of additional proximity calculations on the downstream clustering result, we used a feature selection approach when exploring this aspect of data structure. However, given the popularity of alternative dimensionality reduction methods in scRNA-seq pipelines, such as PCA[62], t-SNE [64] and UMAP [66], it would be beneficial to expand our framework to include approaches based on feature transformation.

Furthermore, t-SNE and UMAP, among many other dimensionality reduction methods, use Euclidean distance as their proximity metric. Given the relatively poor performance of this metric, the application of our framework to explore the influence of other proximity metrics on dimensionality reduction performance may prove insightful [81,82]. Adaptions of the framework could be made to enable evaluation of different data types such as scATAC-seq and DNA methylation. Whilst we applied the same processing pipeline to all of the datasets used in this study, we expect proximity metric performance to be impacted to some extent by dataset processing. As such, we would encourage the design of our framework to be modified to evaluate the potential influence from upstream data handling practices as well.

Taken together, our findings demonstrate how the inherent structural properties of scRNA-seq data have a substantial influence on the performance of proximity measures and subsequently, cell-type clustering and subsequent identification. Given the complexity of scRNA-seq datasets as outlined, it is unlikely for a single metric to perform best in all situations. Instead, we have provided practical guidelines for the selection of proximity metrics likely to perform well with respect to specific properties of the dataset. Furthermore, we provide our framework in the form of a python package to allow users to evaluate proximity metrics for their own datasets. The relevance of this study extends beyond cell clustering, to the numerous scRNA-seq processing and analysis methods which make use of cell-to-cell distances. We hope that the findings from our study and our analysis framework will contribute to improvements in the development of novel metrics and approaches for high-dimensional, sparse data such as scRNA-seq in the future.

## Key Points

- We developed a framework to systematically evaluate the influence of fundamental structural properties of scRNA-seq data on the clustering performance of a diverse range of proximity metrics.
- Clustering performance can be improved substantially by the selection of an appropriate proximity metric and neighbourhood size for the structural properties of a given dataset, and we provide readers with practical guidelines to facilitate this process.
- Many of the proximity metrics’ clustering performance was improved by reduction of dataset sparsity and/or dimensionality.
- Popular metrics such as Euclidean and Manhattan distance performed poorly in comparison to several lessor applied metrics including Cosine, Bray-Curtis and Pearson and Kendall correlations.
- Clustering accuracy with respect to rare cell populations cannot be effectively evaluated by metrics such as ARI and AMI due to their sensitivity to cluster size, and we recommend inclusion of size-independent metrics such as the Pair Sets Index in future studies for situations where bias towards clusters of different sizes is not useful.

## Data Availability

The CellSIUS dataset used to represent discretely structured scRNA-seq data in this article was produced by Wegmann *et al.* [30] and is available in Zenodo: https://zenodo.org/record/3238275. The Fetal Liver Haematopoiesis dataset used to represent continuously structured scRNA-seq data was produced by Popescu *et al.* [43] and is available from the Developmental Human Cell Atlas: https://developmentcellatlas.ncl.ac.uk/datasets/hca_liver/data_share/. Our results, along with raw and processed copies of the datasets used in this study, including simulated scRNA-seq and subsets generated from CellSIUS and the Fetal Liver Haematopoiesis datasets, are available at https://doi.org/10.5281/zenodo.6443267. The evaluation framework package scProximitE along with code to reproduce all figures is available at https://github.com/Ebony-Watson/scProximitE.

## Author Contributions

E.R.W., A.T.F. and J.C.M. formulated the problem. E.R.W developed the evaluation framework and software with input from A.M., J.C.M. and A.T.F. E.R.W designed and implemented the simulations and applied the framework to the case studies on real scRNA-seq data with assistance from A.M. E.R.W., A.T.F. and J.C.M. interpreted the results with input from A.M. E.R.W. wrote the manuscript with input from A.T.F and J.C.M. All authors read and approved the final version of the manuscript.

