## Supplementary Material for "How does data structure impact cell-cell similarity? Evaluating the influence of structural properties on proximity metric performance in single cell RNA-seq data"

#### Supplementary Figures and Tables

### Supplementary Figures:

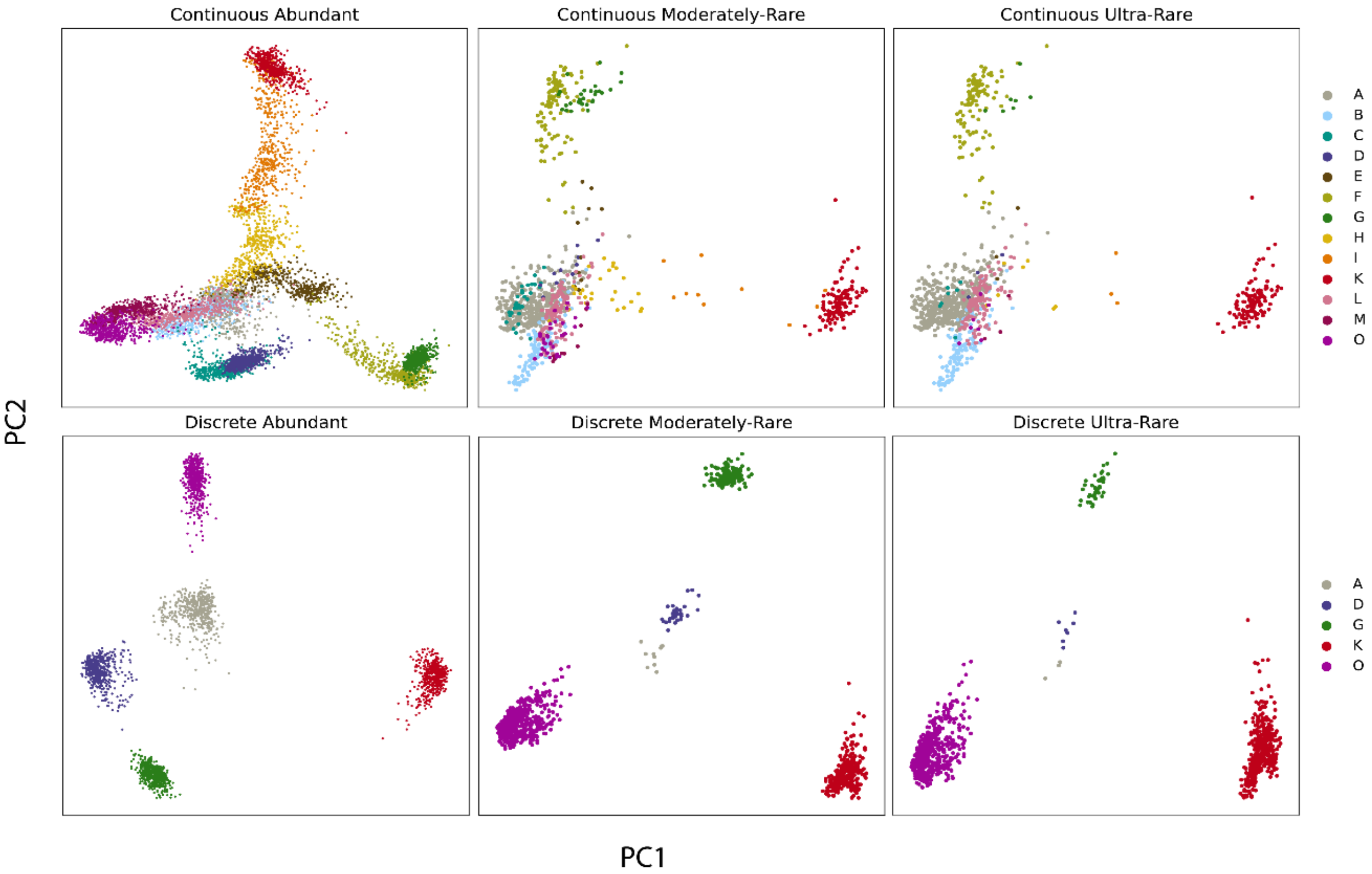


**Supplementary Figure 1:** Principal Components Analysis (PCA) on normalised and scaled data from the Continuous (top) and Discrete (bottom) simulated scRNA-seq data structures. Rows show the Abundant (left), Moderately-Rare (centre), and Ultra-Rare (right) subsets of each structure. Cells are coloured by branch segment, representing the individual cell populations to be identified.

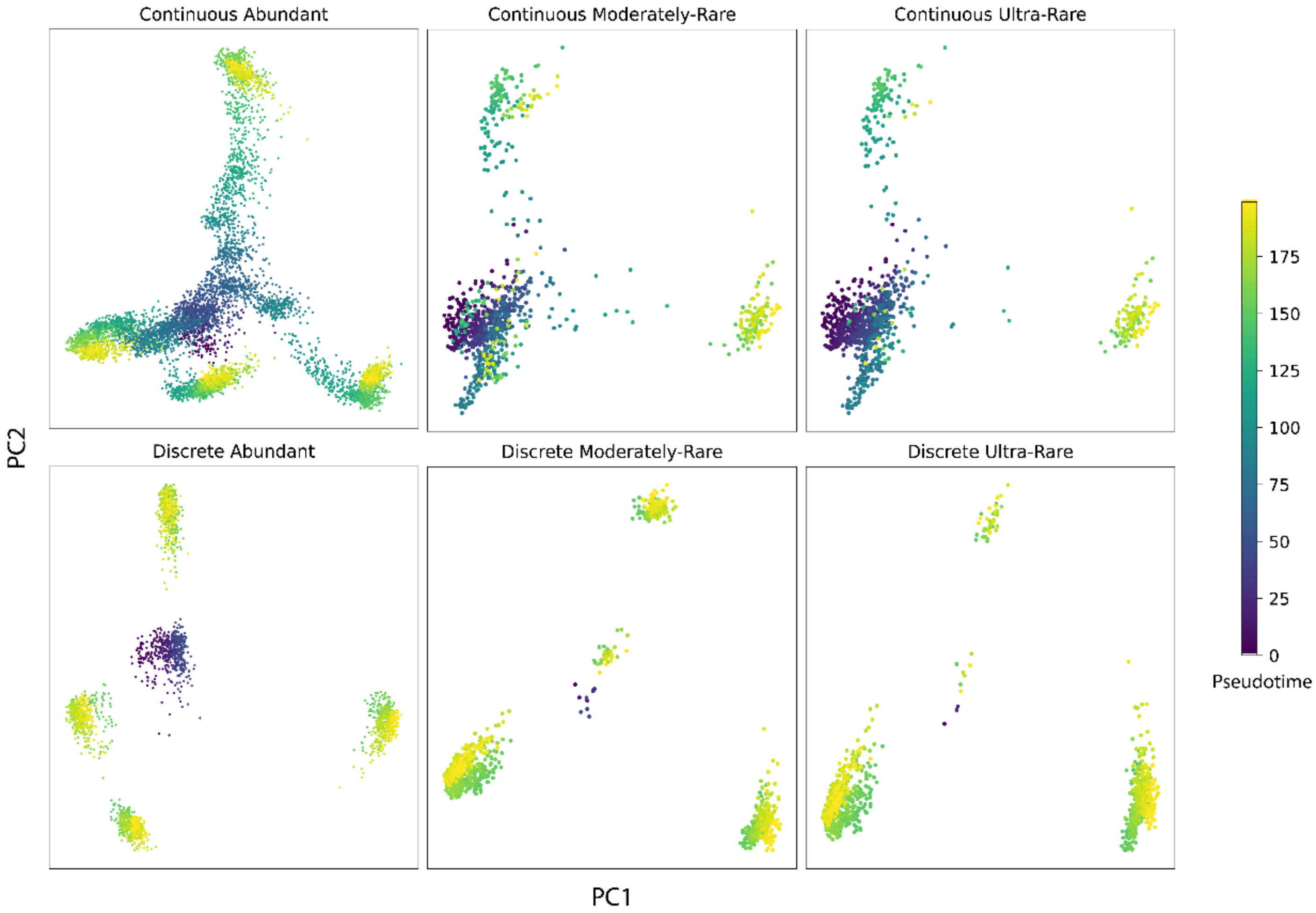


**Supplementary Figure 2:** Principal Components Analysis (PCA) on normalised and scaled data from the Continuous (top) and Discrete (bottom) simulated scRNA-seq data structures. Rows show the Abundant (left), Moderately-Rare (centre), and Ultra-Rare (right) subsets of each structure. Cells are coloured by pseudo-time, representing the distance in pseudo-time units of each cell from the origin (0).

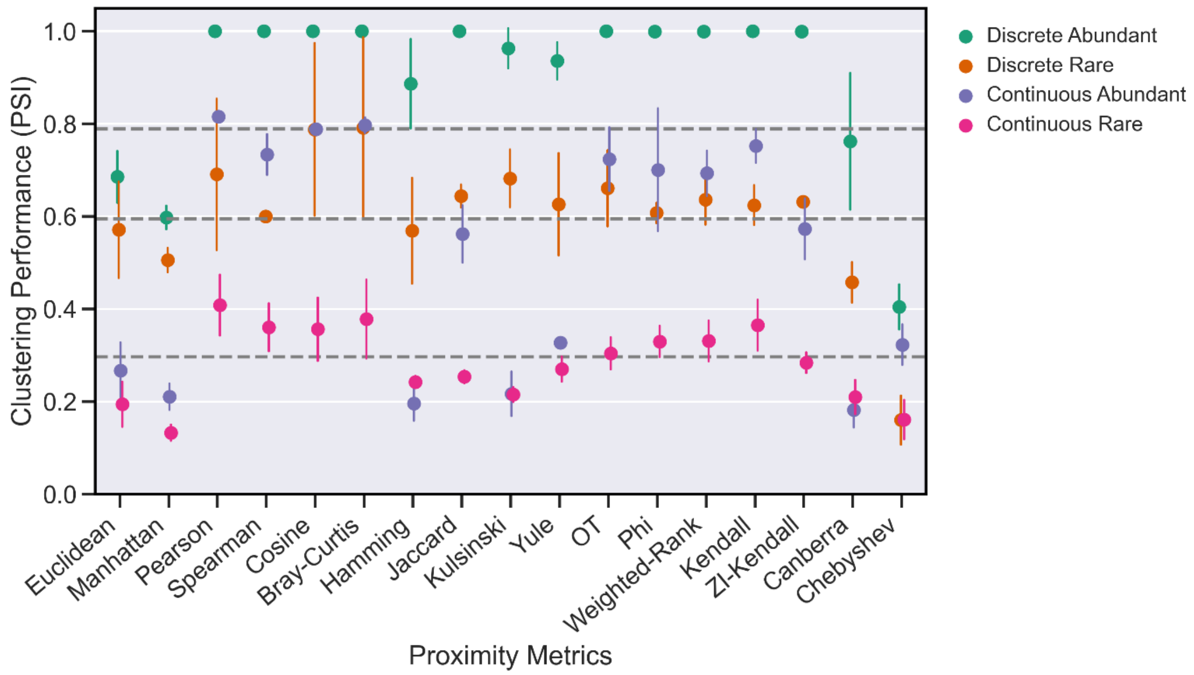


**Supplementary Figure 3:** Clustering performance of proximity metrics for the simulated scRNA-seq datasets (moderate sparsity) representing the four classes of data structure: Discrete Abundant, Discrete Rare, Continuous Abundant, Continuous Rare. Points depict the mean Pair Sets Index (PSI) of clustering from neighbourhood sizes of $k = (3,10,30,50)$, with error bars depicting one standard deviation. Horizontal lines depict (top to bottom) 75th, 50th and 25th percentiles.

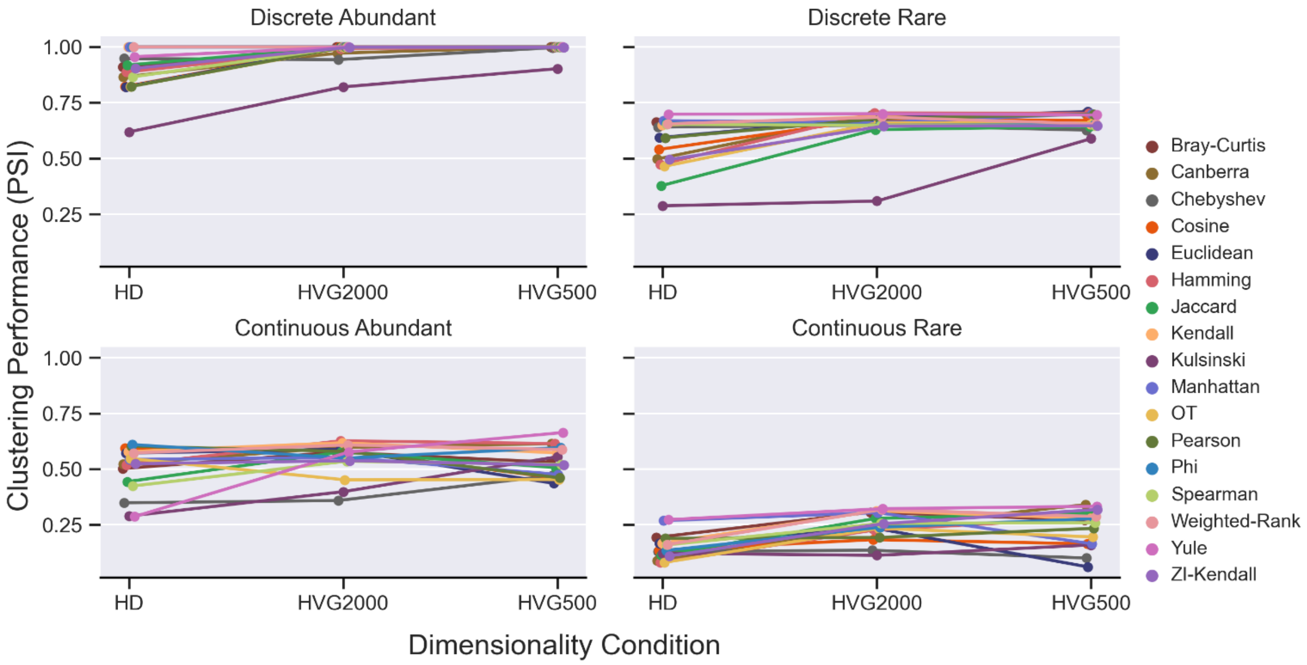


**Supplementary Figure 4**: Clustering performance of all 17 proximity metrics for each structural condition at high-dimensionality (HD) and two levels of reduced dimensionality based on selection of the top 2000 (HVG2000) and 500 (HVG500) highly variable genes. Points depict the mean Pair Sets Index (PSI) of clustering from neighbourhood sizes of $k = (3,10,30,50)$.

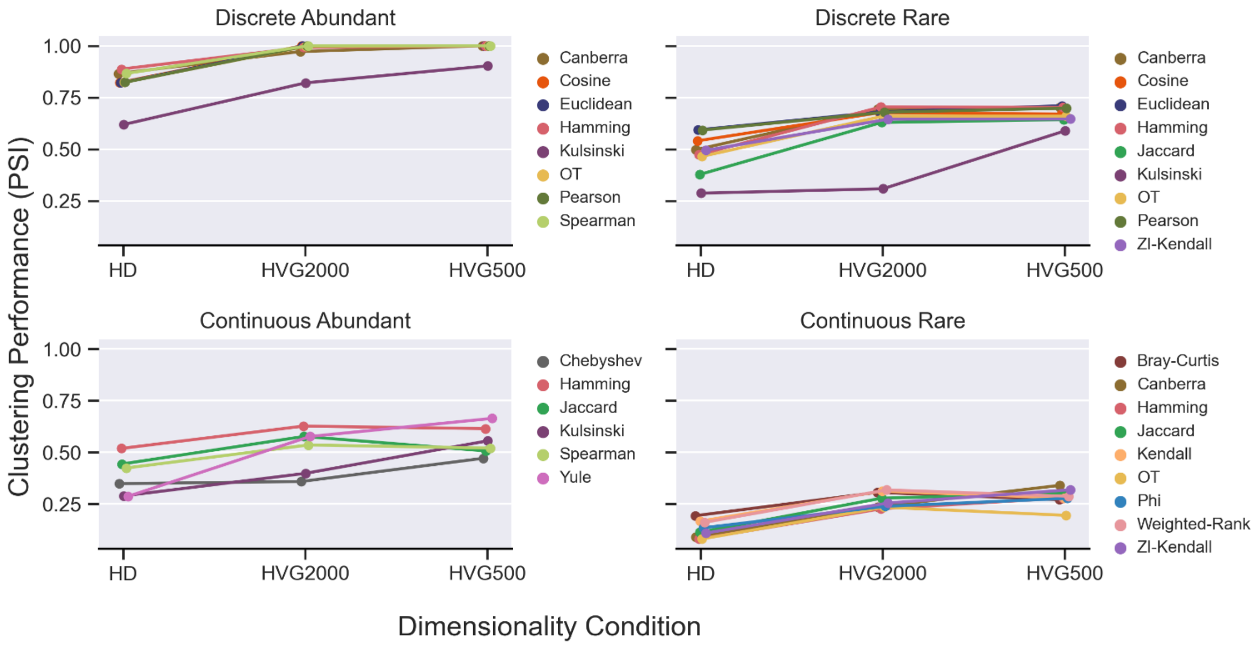


**Supplementary Figure 5:** Clustering performance of the proximity metrics with a >0.1 increase in Pair Sets Index (PSI) change between high-dimensional (HD) dataset and either level of dimensionality reduction (HVG2000, HVG500), for each structural condition. Points depict the mean Pair Sets Index (PSI) of clustering from neighbourhood sizes of $k$ = (3,10,30,50).

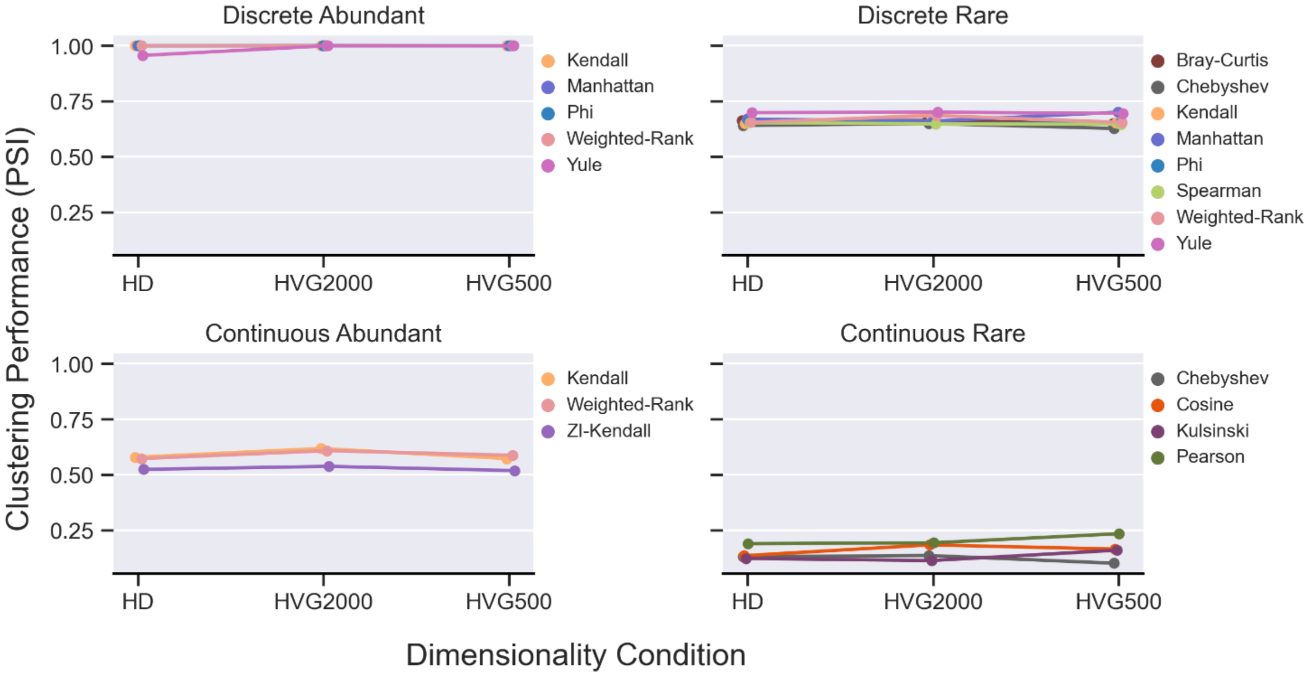


**Supplementary Figure 6**: Performance of all proximity metrics which were identified as invariant, with <0.05 change in between high-dimensional (HD) dataset and either level of dimensionality reduction (HVG2000, HVG500), for each structural condition. Points depict the mean Pair Sets Index (PSI) of clustering from neighbourhood sizes of $k$ = (3,10,30,50).

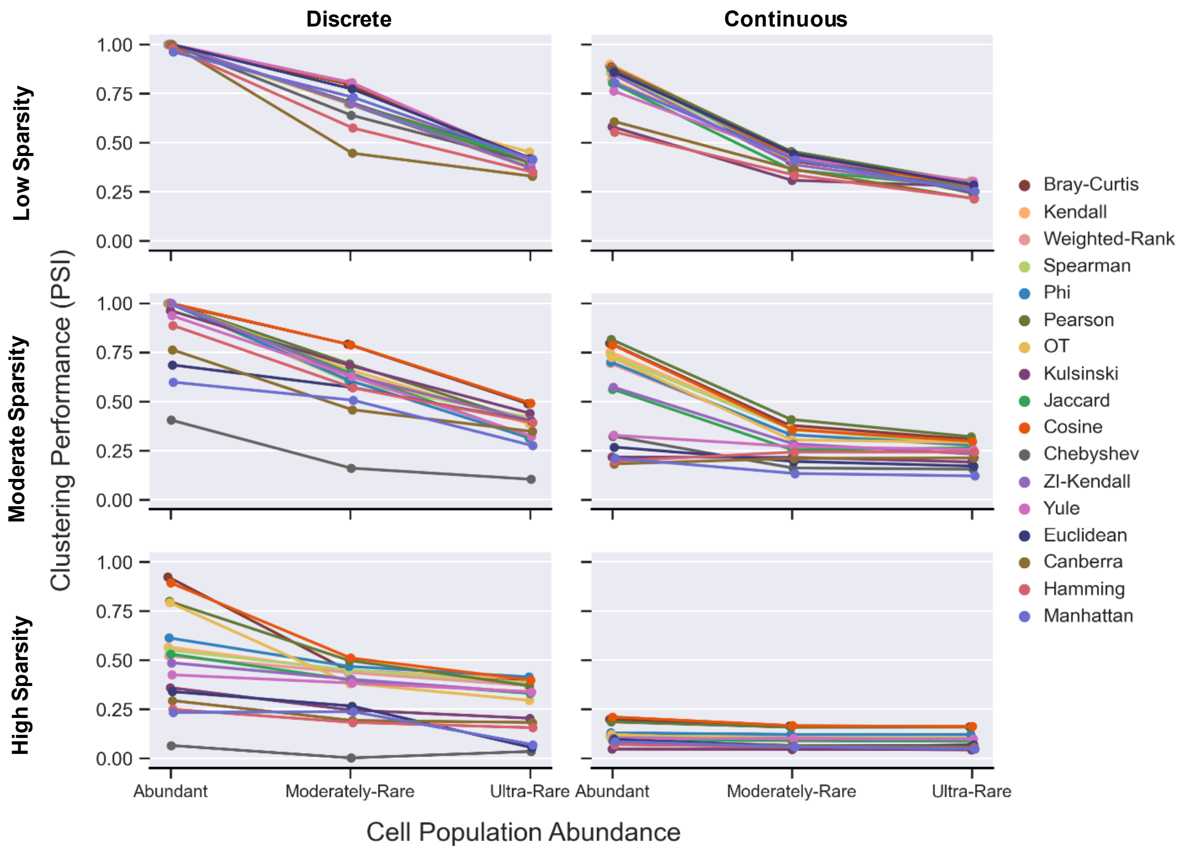


**Supplementary Figure 7**: Clustering performance of all 17 proximity metrics, for Abundant, Moderately-Rare, and Ultra-Rare subsets for the Discrete (left) and Continuous (right) simulated scRNA-seq datasets (moderate sparsity). Points depict the mean Pair Sets Index (PSI) of clustering from neighbourhood sizes of $k = (3,10,30,50)$.

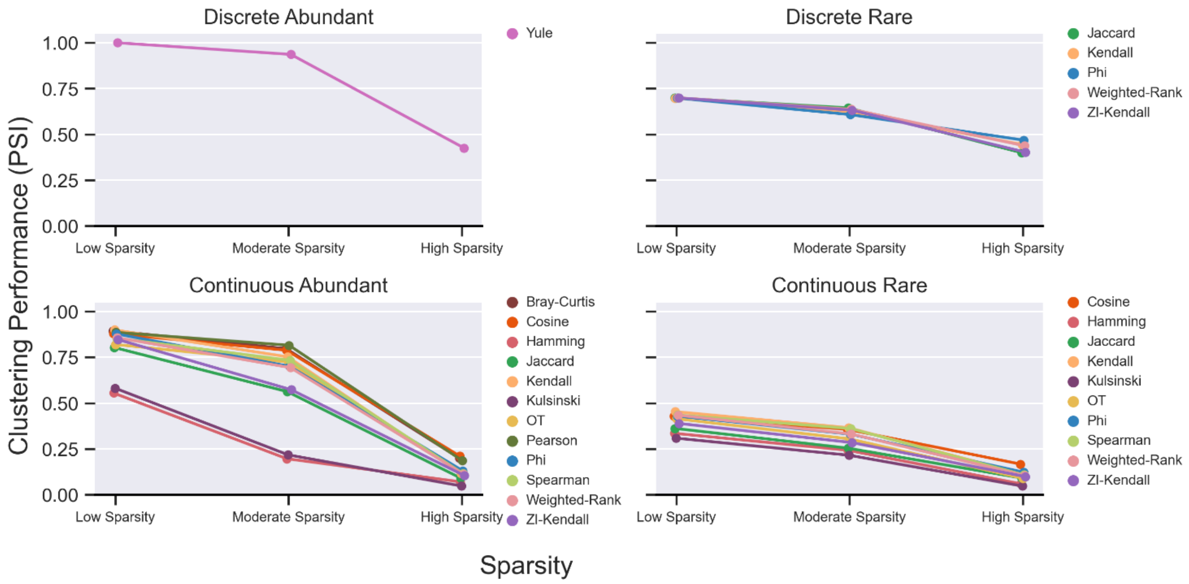


**Supplementary Figure 8**: Performance of proximity metrics identified as moderately sensitive between low (50%) and moderate (70%) sparsity, given a threshold of ≥0.05 but <75th percentile change in PSI. Points depict the mean Pair Sets Index (PSI) of clustering from neighbourhood sizes of $k = (3,10,30,50)$.

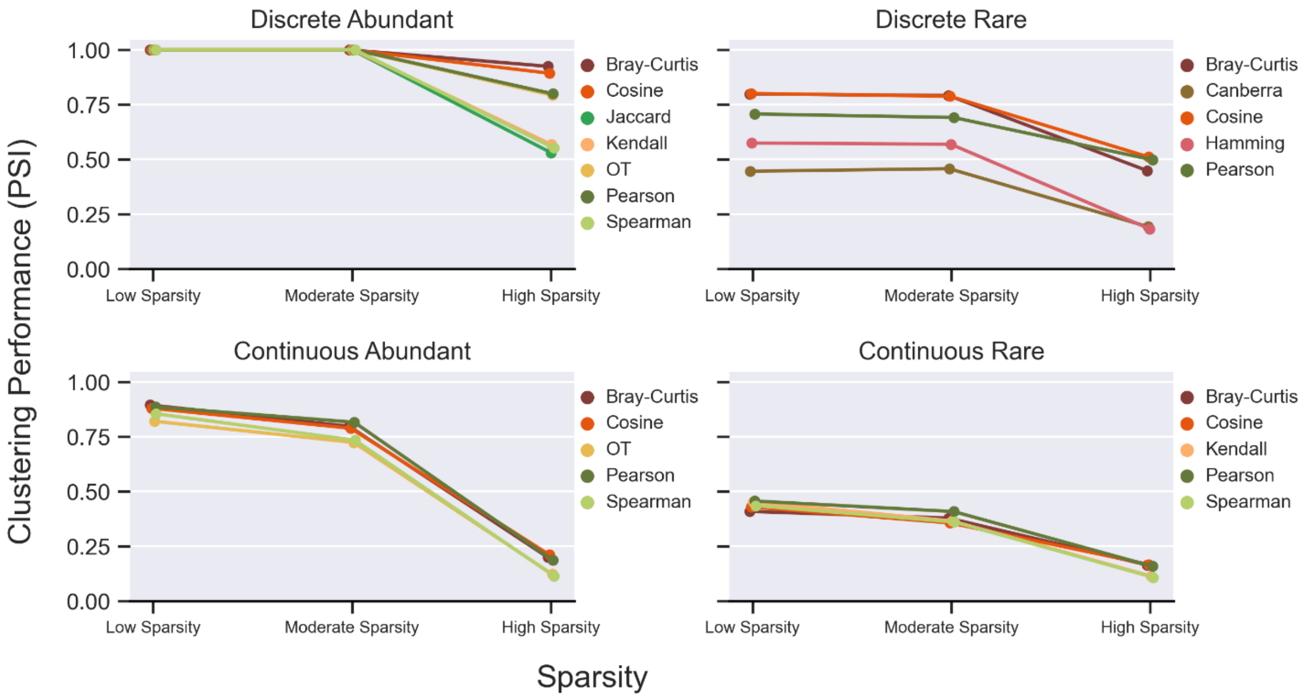


**Supplementary Figure 9**: Performance of the top 5 proximity metrics identified as ranked by smallest change in PSI between low (50%) and moderate (70%) sparsity. Points depict the mean Pair Sets Index (PSI) of clustering from neighbourhood sizes of $k = (3,10,30,50)$.

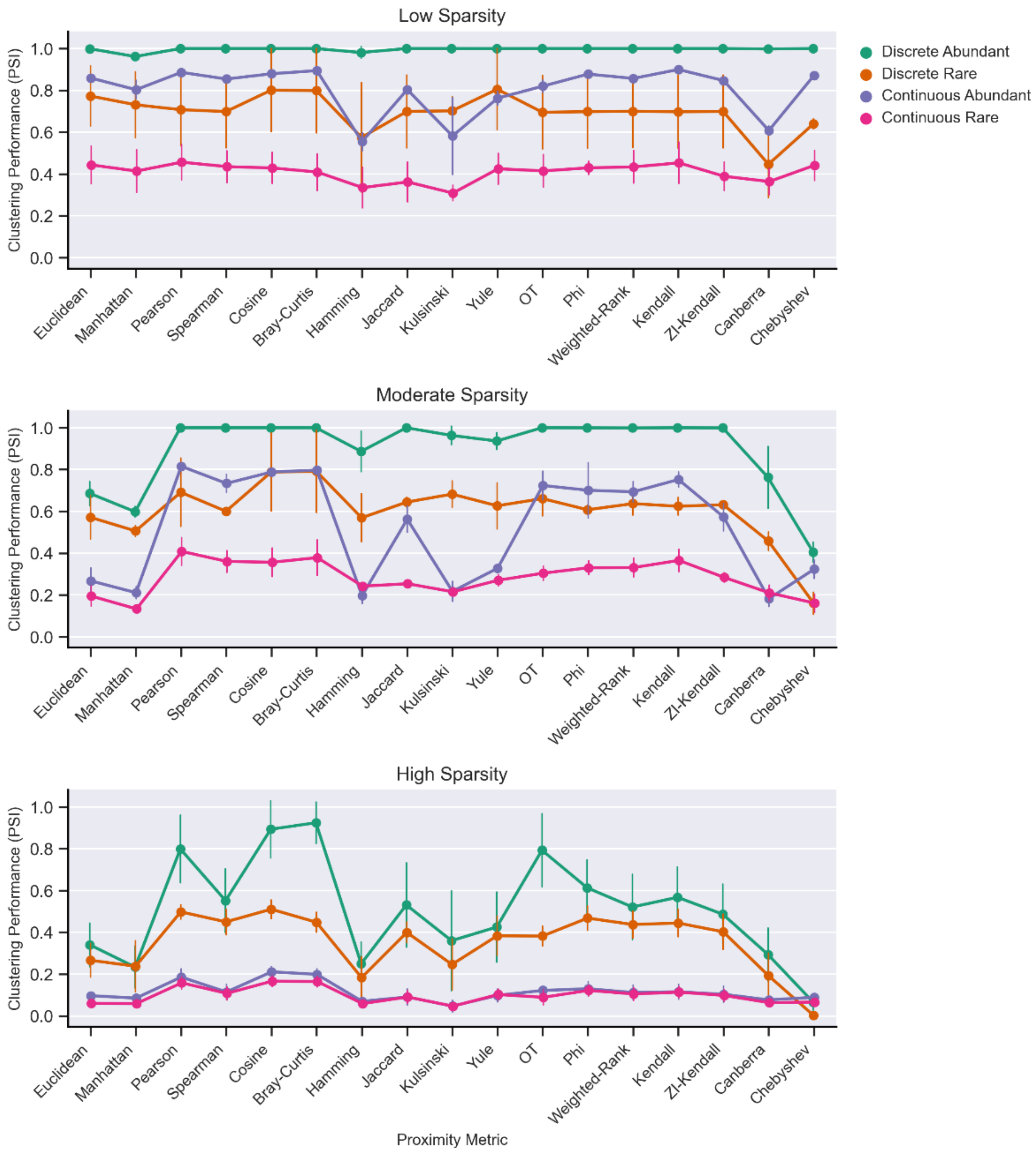


**Supplementary Figure 10**: Clustering performance of all 17 proximity metrics at low (top), moderate (middle) and high (bottom) levels of sparsity, for each structural condition. Points depict the mean Pair Sets Index (PSI) of clustering from neighbourhood sizes of $k$ = (3,10,30,50), with error bars depicting one standard deviation.

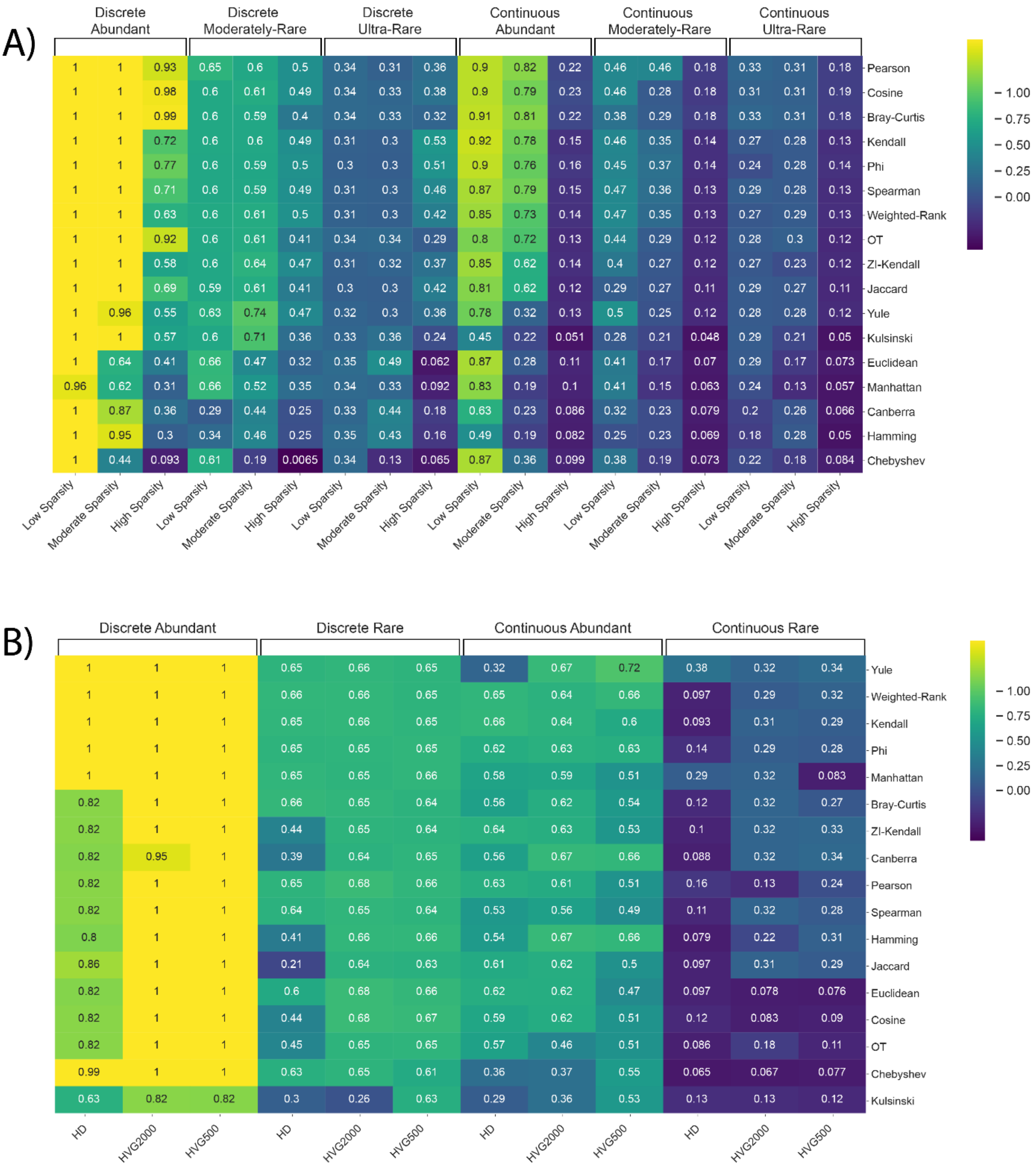


**Supplementary Figure 11**: Metric performance across real scRNA-seq datasets of varying structure and **A)** sparsity and **B)** dimensionality. Heatmap cells contain maximum Pair Sets Index (PSI) obtained at a neighbourhood size $k = 30$, for each metric and dataset combination. Heatmap rows are ordered by mean PSI across datasets.

### Supplementary Tables

**Supplementary Table 1:** Breakdown of cell number and proportion of sub-populations for the CellSIUS [1,2] and the Fetal Liver Haematopoiesis [3,4] datasets.

| Discrete Structure – CellSIUS | | | | | Continuous Structure - Fetal Liver Haematopoiesis | | | | |
|---|---|---|---|---|---|---|---|---|---|
| Cell Line | Abundant | | Rare | | Cell type | Abundant | | Rare | |
| | Cell Number | % | Cell Number | % | | Cell Number | % | Cell Number | % |
| A549 | 400 | 8.0 | 80 | 2.0 | HSC/MPP[1] | 1000 | 20 | 100 | 2.5 |
| H1437 | 270 | 5.4 | 3 | 0.08 | MEMP[2] | 1000 | 20 | 3 | 0.075 |
| HCT116 | 1400 | 28.0 | 1599 | 40.1 | Early-erythroid | 1000 | 20 | 2200 | 55 |
| HEK293 | 1600 | 32.0 | 2000 | 50.2 | Mid-erythroid | 1000 | 20 | 1680 | 42 |
| IMR90 | 500 | 10.0 | 100 | 2.5 | Late-erythroid | 1000 | 20 | 17 | 0.425 |
| Jurkat | 100 | 2.0 | 6 | 0.15 | Total | 5000 | 100 | 4000 | 100 |
| K562 | 379 | 7.6 | 70 | 1.8 | | | | | |
| Ramos | 350 | 7.0 | 125 | 3.1 | | | | | |
| Total | 4999 | 100 | 3983 | 100 | | | | | |

[1]Hematopoietic Stem Cell and Multipotent Progenitor

[2]Mega-karyocyte–Erythroid–Mast cell Progenitor

**Supplementary Table 2:** Breakdown of cell number and proportion of sub-populations for the simulated scRNA-seq datasets [5] of Discrete and Continuous structure, subset for Abundant, Moderately-Rare and Ultra-Rare populations. T = Trajectory of Differentiation, B = branch segment of the differentiation trajectory.

| Discrete – Simulated data | | | | | | |
|---|---|---|---|---|---|---|
| Cell-type | Abundant | | Moderately-Rare | | Ultra-Rare | |
| | Cell *n* | % | Cell *n* | % | Cell *n* | % |
| Origin | 500 | 20 | 10 | 1 | 3 | 0.3 |
| T1, B3 | 500 | 20 | 30 | 3 | 7 | 0.7 |
| T2, B3 | 500 | 20 | 160 | 16 | 40 | 4 |
| T3, B3 | 500 | 20 | 300 | 30 | 450 | 45 |
| T4, B3 | 500 | 20 | 500 | 50 | 500 | 50 |
| *Total* | 2500 | 100 | 1000 | 100 | 1000 | 100 |

| Continuous - Simulated data | | | | | | |
|---|---|---|---|---|---|---|
| Cell-type | Abundant | | Moderately-Rare | | Ultra-Rare | |
| | Cell *n* | % | Cell *n* | % | Cell *n* | % |
| Origin | 500 | 7.69 | 400 | 40 | 500 | 50 |
| T1 B1 | 500 | 7.69 | 110 | 11 | 115 | 11.5 |
| T1, B2 | 500 | 7.69 | 30 | 3 | 7 | 0.7 |
| T1, B3 | 500 | 7.69 | 10 | 1 | 3 | 0.3 |
| T2, B1 | 500 | 7.69 | 10 | 1 | 3 | 11.5 |
| T2, B2 | 500 | 7.69 | 110 | 11 | 115 | 0.7 |
| T2, B3 | 500 | 7.69 | 30 | 3 | 7 | 0.3 |
| T3, B1 | 500 | 7.69 | 30 | 3 | 7 | 11.5 |
| T3, B2 | 500 | 7.69 | 10 | 1 | 3 | 0.7 |
| T3, B3 | 500 | 7.69 | 110 | 11 | 115 | 0.3 |
| T3, B1 | 500 | 7.69 | 110 | 11 | 115 | 11.5 |
| T3, B2 | 500 | 7.69 | 10 | 1 | 3 | 0.7 |
| T3, B3 | 500 | 7.69 | 30 | 3 | 7 | 0.3 |
| *Total* | 6500 | 100 | 1000 | 100 | 1000 | 100 |

**Supplementary Table 3:** Details of percentage of sparsity and cell and gene number for each of the simulated scRNA-seq datasets, Pre- and Post-processing.

| **Sparsity:** | Low Sparsity | | | | Moderate Sparsity | | | High Sparsity | | |
|---|---|---|---|---|---|---|---|---|---|---|
| **Processing:** | Pre | | Post | | Pre | Post | | Pre | Post | |
| **Condition** | Cell x Gene | % | % | Cell x Gene | % | % | Cell x Gene | % | % | Cell x Gene |
| Discrete Abundant | 2500 × 5000 | 49.8 | 46.8 | 2500 × 4691 | 71 | 67.8 | 2500 × 4691 | 90 | 85.2 | 2474 × 2412 |
| Discrete Moderately-Rare | 1000 × 5000 | 49.4 | 45.6 | 1000 × 4622 | 69.9 | 67.1 | 1000 × 4622 | 89.9 | 85 | 991 × 2473 |
| Discrete Ultra-Rare | 1000 × 5000 | 48.8 | 44.7 | 1000 × 4593 | 70.5 | 66.4 | 1000 × 4593 | 89.8 | 84.8 | 992 × 2447 |
| Continuous Abundant | 6500 × 5000 | 48.3 | 45 | 6500 × 4669 | 70.1 | 66.7 | 6500 × 4669 | 89.7 | 85 | 6452 × 2537 |
| Continuous Moderately-Rare | 1000 × 5000 | 46.5 | 43.5 | 1000 × 4713 | 69.1 | 65.9 | 1000 × 4713 | 89.3 | 85 | 990 × 2700 |
| Continuous Ultra-Rare | 1000 × 5000 | 46.1 | 43.1 | 1000 × 4714 | 68.9 | 65.7 | 1000 × 4714 | 68.8 | 84.9 | 991 × 2703 |

**Supplementary Table 4**: Cell number, gene number and dataset sparsity before and after processing for the CellSIUS [1,2] and the Fetal Liver Haematopoiesis (FLH) [3,4] datasets.

| Dataset | Structural Condition | Pre-Processing | | Post-Processing | |
|---|---|---|---|---|---|
| | | Cell x Gene | Sparsity (%) | Cell x Gene | Sparsity (%) |
| CellSIUS | Discrete Abundant | 5000 × 23848 | 84.9 | 5000 × 8982 | 63.1 |
| CellSIUS | Discrete Rare | 3984 × 23848 | 85.1 | 3984 × 8802 | 63.1 |
| FLH | Continuous Abundant | 5000 × 27080 | 87.8 | 5000 × 8174 | 62.4 |
| FLH | Continuous Rare | 4000 × 27080 | 85.1 | 4000 × 8622 | 55.4 |

**Supplementary Table 5:** Implementation details for the 17 proximity metrics investigated. All data underwent filtering and normalisation prior to calculation of proximity metric matrices. If the output from a proximity metric function was not in the form of a dissimilarity metric, additional computations were performed to convert them.

| Metric | Class | Input | Language | Package |
|---|---|---|---|---|
| Euclidean | Distance | Counts | Python (v3.8) [6] | sklearn (v1.0.1) [7] |
| Manhattan | Distance | Counts | Python (v3.8) [6] | sklearn (v1.0.1) [7] |
| Canberra | Distance | Counts | Python (v3.8) [6] | SciPy (v1.7.1)[8] |
| Chebyshev | Distance | Counts | Python (v3.8) [6] | SciPy (v1.7.1)[8] |
| Hamming | Correlation | Counts | Python (v3.8) [6] | SciPy (v1.7.1)[8] |
| Pearson | Correlation | Counts | Python (v3.8) [6] | SciPy (v1.7.1)[8] |
| Spearman[1] | Correlation | Counts | Python (v3.8) [6] | SciPy (v1.7.1)[8] |
| Weighted-Rank[1] | Correlation | Counts | R (v4.1.1) [9] | dismay [10] |
| Kendall[1] | Correlation | Counts | R (v4.1.1) [9] | dismay [10] |
| ZI-Kendall[1,2] | Correlation | Counts | R (v4.1.1) [9] | dismay [10] |
| Bray-Curtis | Proportionality | Counts | Python (v3.8) [6] | SciPy (v1.7.1)[8] |
| Cosine | Similarity | Counts | Python (v3.8) [6] | SciPy (v1.7.1)[8] |
| Phi[3] | Proportionality | Counts | R (v4.1.1) [9] | dismay [10] |
| Optimal Transport[4] | Dissimilarity | Counts | Python (v3.8) [6] | Otscomics (v0.0.1)[11] |
| Jaccard index | Dissimilarity | Binarised | Python (v3.8) [6] | SciPy (v1.7.1)[8] |
| Kulsinski | Dissimilarity | Binarised | Python (v3.8) [6] | SciPy (v1.7.1)[8] |
| Yule | Dissimilarity | Binarised | Python (v3.8) [6] | SciPy (v1.7.1)[8] |

[1]To transform the outputted cellxcell matrix values into the form of a dissimilarity, all matrix values were subtracted from 1.

[2]Rescaled outputted cellxcell matrix to contain values in a bounded range (0 – 1) and enforced a 0 diagonal.

[3]To transform the outputted cellxcell matrix values into the form of a dissimilarity, the absolute values of outputted cellxcell matrix were taken.

[4]Requires GPU.

**Supplementary Table 6**: Representative datasets and their respective performance ranges for the recommended proximity metrics and neighbourhood sizes (*k*) structural properties (Figure 9). As simulated scRNA-seq data was used to evaluate the influence of sparsity, these datasets generally contain lower levels of noise and complexity as real scRNA-seq data. As such, higher PSI ranges are typically observed for the simulated datasets.

| Structure | Dataset | Properties | Proximity Measures | *k* | PSI[2] range |
|---|---|---|---|---|---|
| Discrete Abundant | CellSIUS[1] | Reduced dimensionality | Any metric but Kulsinski | Any | PSI >0.99 |
| | Simulated[5] | High Sparsity | Bray-Curtis, Cosine | 30,50,100 | PSI > 0.98 |
| | | Low/Moderate Sparsity | Weighted Rank, Kendall, Phi | Any | PSI >0.99 |
| Discrete Rare | CellSIUS[1] | Reduced dimensionality | Euclidean, Manhattan, Yule, Hamming, Canberra, Pearson, Cosine | 3 | 0.67 < PSI < 0.71 |
| | Simulated[5] | High Sparsity | Phi, Spearman, Kendall | 100 | 0.54 < PSI < 0.64 |
| | | | Pearson, Cosine | 3 | |
| | | Low/Moderate Sparsity | Bray-Curtis, Pearson, Cosine | 3 | 0.67 < PSI < 0.71 |
| Continuous Abundant | FLH[1][4] | Reduced dimensionality | Yule, Kendall, Weighted-Rank | 3,10 | 0.28 < PSI < 0.34 |
| | Simulation[5] | High Sparsity | Bray-Curtis, Pearson, Cosine | 30,50,100 | 0.18 < PSI < 0.2 |
| | | Low/Moderate Sparsity | Bray-Curtis, Kendall, Pearson | 3,10 | 0.41 < PSI < 0.48 |
| Continuous Rare | FLH[1][4] | Reduced dimensionality | Yule, Hamming, Canberra, Kendall, Weighted-Rank | Any | 0.57 < PSI < 0.67 |
| | Simulated[5] | High Sparsity | Bray-Curtis, Pearson, Cosine | 30,50,100 | 0.18 < PSI <0.24 |
| | | Low/Moderate Sparsity | Bray-Curtis, Pearson, Cosine | 30,50,100 | 0.76 < PSI < 0.83 |

[1] Fetal Liver Haematopoiesis.

[2] Pair Sets Index.

1. Wegmann R, Neri M, Schuierer S, et al. CellSIUS provides sensitive and specific detection of rare cell populations from complex single-cell RNA-seq data. Genome Biology 2019; 20:142
2. Wegmann R, Neri M. CellSIUS provides sensitive and specific detection of rare cell populations from complex single cell RNA-seq data: Codes and processed data. 2019;
3. Popescu D-M, Botting RA, Stephenson E, et al. Decoding human fetal liver haematopoiesis: Dataset. 2019;
4. Popescu D-M, Botting RA, Stephenson E, et al. Decoding human fetal liver haematopoiesis. Nature 2019; 574:365–371
5. Papadopoulos N, Gonzalo PR, Söding J. PROSSTT: probabilistic simulation of single-cell RNA-seq data for complex differentiation processes. Bioinformatics 2019; 35:3517–3519
6. Van Rossum G, Drake FL. Python 3 Reference Manual. 2009;
7. Pedregosa F, Varoquaux G, Gramfort A, et al. Scikit-learn: Machine Learning in Python. Journal of Machine Learning Research 2011; 12:2825–2830
8. Virtanen P, Gommers R, Oliphant TE, et al. SciPy 1.0: fundamental algorithms for scientific computing in Python. Nat Methods 2020; 17:261–272
9. R Core Team. R: The R Project for Statistical Computing. 2021;
10. Skinnider MA, Squair JW, Foster LJ. Evaluating measures of association for single-cell transcriptomics. Nat Methods 2019; 16:381–386
11. Huizing G-J, Peyré G, Cantini L. Optimal Transport improves cell-cell similarity inference in single-cell omics data. 2021; 2021.03.19.436159
